# Calibrating the Human Mutation Rate via Ancestral Recombination Density in Diploid Genomes

**DOI:** 10.1101/015560

**Authors:** Mark Lipson, Po-Ru Loh, Sriram Sankararaman, Nick Patterson, Bonnie Berger, David Reich

## Abstract

The human mutation rate is an essential parameter for studying the evolution of our species, interpreting present-day genetic variation, and understanding the incidence of genetic disease. Nevertheless, our current estimates of the rate are uncertain. Most notably, recent approaches based on counting *de novo* mutations in family pedigrees have yielded significantly smaller values than classical methods based on sequence divergence. Here, we propose a new method that uses the fine-scale human recombination map to calibrate the rate of accumulation of mutations. By comparing local heterozygosity levels in diploid genomes to the genetic distance scale over which these levels change, we are able to estimate a long-term mutation rate averaged over hundreds or thousands of generations. We infer a rate of 1.61 ± 0.13 × 10^−8^ mutations per base per generation, which falls in between phylogenetic and pedigree-based estimates, and we suggest possible mechanisms to reconcile our estimate with previous studies. Our results support intermediate-age divergences among human populations and between humans and other great apes.

## Author Summary

The rate at which new heritable mutations occur in the human genome is a fundamental parameter in population and evolutionary genetics. However, recent direct family-based estimates of the mutation rate have consistently been much lower than previous results from comparisons with other great ape species. Because split times of species and populations estimated from genetic data are often inversely proportional to the mutation rate, resolving the disagreement would have important implications for understanding human evolution. In our work, we apply a new technique that uses mutations that have accumulated over many generations on either copy of a chromosome in an individual’s genome. Instead of an external reference point, we rely on fine-scale knowledge of the human recombination rate to calibrate the long-term mutation rate. Our procedure accounts for possible errors found in real data, and we also show that it is robust to a range of model violations. Using eight diploid genomes from non-African individuals, we infer a rate of 1.61 ± 0.13 × 10^−8^ single-nucleotide changes per base per generation, which is intermediate between most phylogenetic and pedigree-based estimates. Thus, our estimate implies reasonable, intermediate-age population split times across a range of time scales.

## Introduction

All genetic variation—the substrate for evolution—is ultimately due to spontaneous heritable mutations in the genomes of individual germline cells. The most commonly studied mutations are point mutations, which consist of single-nucleotide changes from one base to another. The rate at which these changes occur, in combination with other forces, determines the frequency with which homologous nucleotides differ from one individual’s genome to another.

A number of different approaches have previously been used to estimate the human mutation rate [1–3], of which we mention four categories here. The first method is to count the number of fixed genetic changes between humans and another species, such as chimpanzees [4]. Population genetic theory implies that if the mutation rate remains constant, then neutral mutations (those that do not affect an organism’s fitness) should accumulate between two genomes at a constant rate (the well-known “molecular clock” [5]). Thus, the mutation rate can be estimated based on the divergence time of the genomes, if this can be confidently inferred from fossil evidence. However, even if the age of fossil remains can be accurately determined, assigning their proper phylogenetic positions is often difficult. Moreover, because of shared ancestral polymorphism, the time to the most recent common ancestor is always older—and sometimes far older—than the time of species divergence, meaning that split-time calibrations cannot always be directly applied to genetic divergences.

A second common approach, which has only become possible within the last few years, is to count newly occurring mutations in deep sequencing data from family pedigrees, especially parent-child trios [6–10]. This approach provides a direct estimate but can be technically challenging, as it is sensitive to genotype accuracy and data processing from high-throughput sequencing. In particular, sporadic sequencing and alignment errors can be difficult to distinguish from true *de novo* mutations. Surprisingly, these sequencing-based estimates have consistently been much lower than those based on the first approach: in the neighborhood of 1−1.2 × 10^−8^ per base per generation, as opposed to 2−2.5 × 10^−8^ for those from long-term divergence [1–3].

A third method, and another that is only now becoming possible, is to make direct comparisons between present-day samples and precisely-dated ancient genomes. This method is similar to the first one, but by using two time-separated samples from the same species, it avoids the difficulty of needing an externally inferred split time. A recent study of a high-coverage genome sequence from a 45,000-year-old Upper Paleolithic modern human produced two estimates of this type [11]. Direct measurement of decreased mutational accumulation in this sample led to rate estimates of 0.44−0.63 × 10^−9^ per base per year (range of 14 estimates), or 1.3−1.8 × 10^−8^ per base per generation (assuming 29 years per generation [12]). An alternative technique, leveraging time shifts in historical population sizes, yielded an estimate of 0.38−0.49 × 10^−9^ per base per year (95% confidence interval), or 1.1−1.4 × 10^−8^ per base per generation, although a re-analysis of different mutational classes led to a total estimate of 0.44−0.59 × 10^−9^ per base per year (1.3−1.7 × 10^−8^), in better agreement with the first approach [11].

Finally, a fourth technique is to calibrate the rate of accumulation of mutations using a separate evolutionary rate that is better measured. In one such study, the authors used a model coupling single-nucleotide mutations to mutations in nearby microsatellite alleles to infer a single-nucleotide rate of 1.4−2.3 × 10^−8^ per base per generation (90% confidence interval) [13]. In principle, this general technique is appealing because it only involves intrinsic information, without any reference points, and yet can leverage the signal of mutations that have occurred over many generations.

In this study, we present a new approach that falls into this fourth category: we calibrate the mutation rate against the rate of meiotic recombination events, which has been measured with high precision in humans [14–17]. Intuitively, our method makes use of the following relationship between the mutation and recombination rates. At every site *i* in a diploid genome, the two copies of the base have some time to most recent common ancestor (TMRCA) *T_i_*, measured in generations. The genome can be divided into blocks of sequence that have been inherited together from the same common ancestor, with different blocks separated by ancestral recombinations. If a given block has a TMRCA of *T* and a length of *L* bases, and if *μ* is the per-generation mutation rate per base, then the expected number of mutations that have accumulated in either copy of that block since the TMRCA is 2*T Lμ.* This is the expected number of heterozygous sites that we observe in the block today (disregarding the possibility of repeat mutations). We also know that if the per-generation recombination rate is *r* per base, then the expected length of the block is (2*Tr*)^−1^. Thus, the expected number of heterozygous sites per block (regardless of age) is *μ/r.*

This relationship allows us to estimate *μ* given a good prior knowledge of *r*. Our full method is more complex but is based on the same principle. We show below how we can capture the signal of heterozygosity per recombination to infer the historical per-generation mutation rate for non-African populations over approximately the last 50–100 thousand years (ky). A broadly similar idea is also applied in an independent study [18], but over a more recent time scale (up to ~ 3 ky, via mutations present in inferred identical-by-descent segments), and the two final estimates are in very good agreement.

## Results

### Overview of methods

One difficulty of the simple method outlined above is that in practice we cannot accurately reconstruct the breakpoints between adjacent non-recombined blocks. Instead, we use an indirect statistic that captures information about the presence of breakpoints but can be computed in a simple way (without directly inferring blocks) and averaged over many loci (Fig 1).

**Figure 1.**
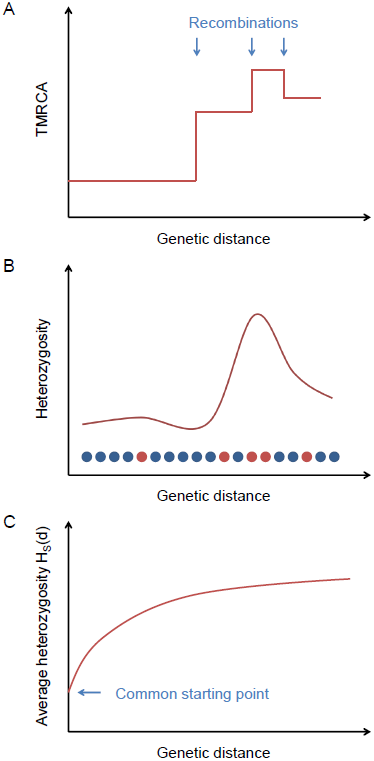
Explanation of the statistic *H_S_*(*d*). (A) Ancestral recombinations separate chromosomes into blocks of piecewise-constant TMRCA (and hence expected heterozygosity). (B) From the data, we measure local heterozygosity as a function of genetic distance; red and blue circles represent heterozygous and homozygous sites, respectively, along a diploid genome. (C) Our statistic *H_S_*(*d*) is an average heterozygosity as a function of genetic distance over many starting points with similar local heterozygosities, yielding a smooth relaxation toward the genome-wide average.

Starting from a certain position in the genome, the TMRCA of the two haploid chromosomes as a function of distance in either direction is a step function, with changes at ancestral recombination points (Fig 1A). Heterozygosity, being proportional to TMRCA in expectation (and directly observable), follows the same pattern on average (Fig 1B).Recombinations

If we consider a collection of starting positions having similar local heterozygosities, then as a function of the genetic distance *d* away from them, the average heterozygosity displays a smooth relaxation from the common starting value toward the global mean heterozygosity 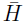, as the probability increases of having encountered recombination points (Fig 1C). We define a statistic *H_S_*(*d*) that equals this average heterozygosity, where *S* is a set of starting points indexed by the local number of heterozygous sites per 100 kb (we also use S at times to refer to the heterozygosity range itself). The TMRCAs of these points determine the time scale over which our inferred value of *μ* is measured. Our default choice is to use starting points with a local total of 5–10 heterozygous sites per 100 kb (see Methods).

To estimate *μ*, we use the fact that the probability of having encountered a recombination as one moves away from a starting point is a function of both *d* and the starting heterozygosity *H_S_*(0), since smaller values of *H_S_*(0) correspond to smaller TMRCAs, with less time for recombination to have occurred, and hence longer unbroken blocks. This relationship allows us to calibrate *μ* against the recombination rate *r* via the relaxation rate of *H_S_*(*d*). Our inference procedure involves using coalescent simulations to create matching “calibration data” with known values of *μ* and then solving for the best-fit mutation rate for the test data (see Methods and Fig 2). We note that when comparing *H_S_*(*d*) for real data to the calibration curves, a larger value of *μ* will correspond to a lower curve. This is because *H_S_*(0) is fixed, which means that the TMRCAs at the starting points are proportionally lower for larger values of *μ*. Thus, recombinations are less frequent as a function of *d*, leading to a slower relaxation.

**Figure 2.**
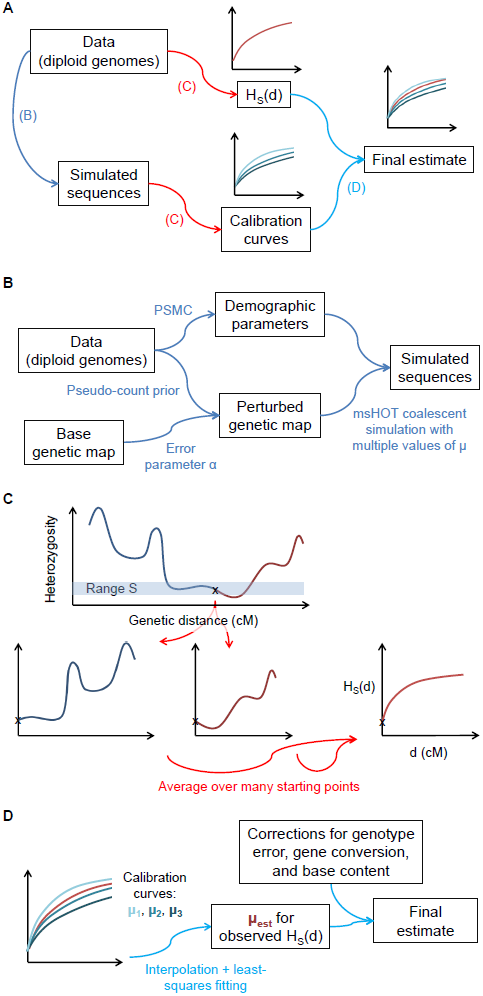
Illustration of the steps of our inference procedure. (A) Overview: from the data, we compute both the statistic *H_S_*(*d*) and other parameters necessary to create matching calibration curves with known values of *μ*. (B) Details of capturing aspects of the real data for the calibration data. (C) Computation of *H_S_*(*d*): the statistic captures the average heterozygosity as a function of genetic distance *d* from a starting point with heterozygosity in a defined range *S*, averaged over many such points. (D) For the final inferred value of *μ*, we compare matched *H_S_*(*d*) curves for the real data and calibration data (with known values of *μ*).

In order for our inferences to be accurate, the calibration curves must recapitulate as closely as possible all aspects of the real data that could affect *H_S_*(*d*) (see Methods, S1 Text, and Fig 2). First, because coalescent probabilities depend on ancestral population sizes, we use PSMC [19] to learn the demographic history of our samples. Next, we adapt a previously developed technique [20] to infer the fine-scale uncertainty of our genetic map. Finally, we correct our raw inferred values of *μ* for three additional factors in order to isolate the desired mutational signal: (1) we multiply by a correction for genotype errors; (2) we subtract the contribution of non-crossover gene conversion, using a result from [21] adjusted for local recombination rate; and (3) we scale the final value to correspond to genome-wide base content and mutability (see Methods and S1 Text). We also test additional potential model violations through simulations (see S1 Text and S1 Fig). We account for statistical uncertainty using a block jackknife and incorporate confidence intervals for model parameters; all results are given as mean ± standard error.

## Simulations

First, for seven different scenarios, including a range of possible model violations, we generated 20 simulated diploid genomes with a known true mutation rate (*μ* = 2.5 × 10^−8^ per generation except where otherwise specified) and ran our procedure as we would for real data, with perturbed genetic maps for both the test data and calibration data (variance parameter *α* = 3000 M^−1^; see Methods). To measure the uncertainty in our estimates, we performed 25 independent trials of each simulation, and we also compared the standard deviations of the estimates across trials with jackknife-based standard errors (as we would measure uncertainty for real data). Full details of the simulation procedures can be found in Methods and S1 Text.

In all cases, the *H*_5−10_(*d*) curves matched quite well between the test data and the calibration data, and our final results were within two standard errors of the true rate (Fig 3). Furthermore, our jackknife estimates of the standard error were comparable to the realized standard deviations and on average conservative, especially for the most complex simulation (g), despite not incorporating PSMC uncertainty (see Methods): 0:08 × 10^−8^, 0:04 × 10^−8^, 0:04 × 10^−8^, 0:06 × 10^−8^, 0:09 × 10^−8^, 0:05 × 10^−8^, and 0:11 × 10^−8^, respectively, for the seven scenarios (see Fig 3 for empirical standard deviations). The fact that all of the inferred rates are close to the true values leads us to conclude that none of the aspects of the basic procedure or the tested model violations create a substantial bias.

**Figure 3.**
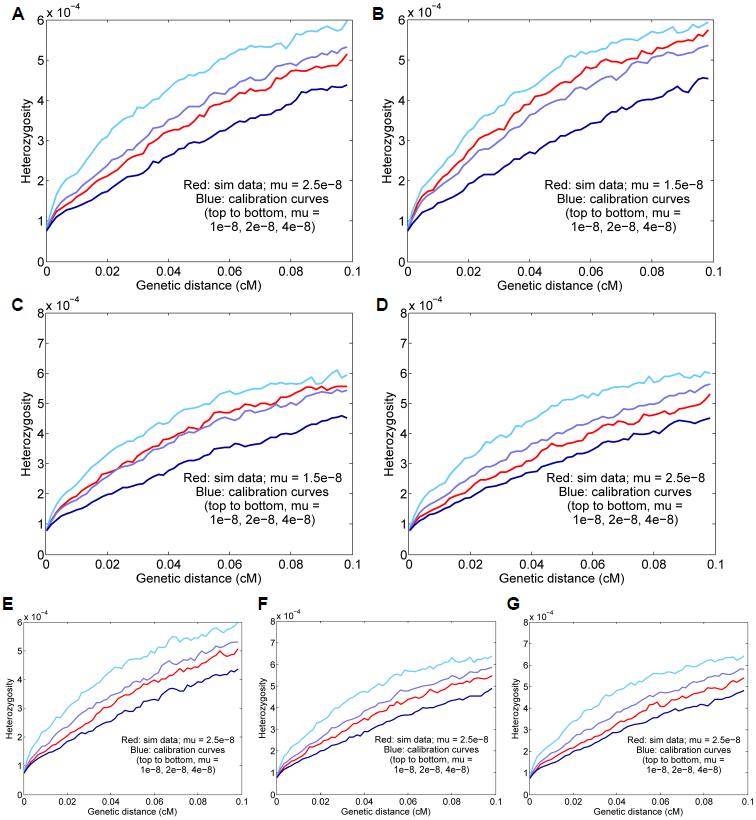
Results for simulated data. Means and standard deviations of 25 independent trials are given, and the curves displayed are for representative runs matching the 25-trial means. The true simulated rate is *μ* = 2.5 × 10^−8^ unless otherwise specified. (A) Baseline simulated data; the inferred rate is *μ* = 2.47 ± 0.05 × 10^−8^. (B) Basic simulated data with a true rate of 1.5 × 10^−8^; the inferred rate is *μ* = 1.57 ± 0.04 × 10^−8^. (C) Data with a true rate of 1.5 × 10^−8^ plus gene conversion; the inferred rate is *μ* = 1.49 ± 0.05 × 10^−8^ (corrected from a raw value of 1.70 × 10^−8^ with gene conversion included). (D) Data with simulated genotype errors; the inferred rate is *μ* = 2.39 ± 0.06 × 10^−8^ (corrected from a raw value of 2.71 × 10^−8^ with genotype errors included). (E) Data simulated with variable mutation rate; the inferred rate is *μ* = 2.61 ± 0.08 × 10^−8^. (F) Data from a simulated admixed population; the inferred rate is *μ* = 2.57 ± 0.07 × 10^−8^. (G) Simulated data with all three complications as in (D)–(F); the inferred rate is *μ* = 2.53 ± 0.06 × 10^−8^ (corrected from a raw value of 2.77 × 10^−8^).

## Error parameters

Before obtaining mutation rate estimates from real data, we quantified two important error parameters: the rate of false heterozygous genotype calls and the degree of inaccuracy in our genetic map.

We estimated the genotype error rate by taking advantage of the fact that methylated cytosines at CpG dinucleotides are roughly an order of magnitude more mutable than other bases [3, 7, 8,10] (see Methods). Thus, such mutations are strongly over-represented among true heterozygous sites as compared to falsely called heterozygous sites. By counting the proportion of CpG mutations out of all heterozygous sites around our ascertained starting points, we inferred an error rate of approximately 1 per 100 kb (1.08 ± 0.28 × 10^−5^ per base; see Methods and S1 Text), consistent with previous results [22].

It was also necessary for us to estimate the accuracy of our genetic map. We used the “shared” version of the African-American (AA) map from [17] as our base map and a modified version of the error model of [20]: *Z* ~ Gamma(*αγ*(*g* + *πp*), *α*), where *Z* is the true genetic length of a map interval, *g* is the observed genetic length, *p* is the physical length, *α* is the parameter measuring the accuracy of the map, and *γ* and *π* are constants (see Methods). Based on pedigree crossover data from [23], we estimated *α* = 2802 ± 14 M^−1^ for the full AA map and *α* = 3414 ± 13 M^−1^ for the “shared” map, which should serve as lower and upper bounds (see Methods). For our analyses, we took *α* = 3100 M^−1^ (with a standard error of 300 M^−1^ to account for our uncertainty in the precise value). This means that 1/*α* ≈ 0.03 cM can be thought of as the length scale for the accuracy of genetic distances according to the base map (see Methods for details). In order to translate the uncertainty in *α* into its effect on the inferred *μ*, we repeated our primary analysis with a range of alternative values of *α* (S2 Fig).

We note that the values of α reported in [20] are substantially lower than ours, which we suspect is because our validation data have much finer resolution than those used previously. (When using the same validation data, the “shared” and HapMap LD [15] maps appear to be relatively similar in accuracy.) If we substitute our new α values for the original application of inferring the date of Neanderthal gene flow into modern humans, we obtain a less distant time in the past, 28–65 ky (most likely 35–49 ky), versus 37–86 ky (most likely 47–65 ky) reported in [20]. While relatively recent, this date range is not in conflict with archaeological evidence or with an estimate of 49–60 ky (95% confidence interval) based on an Upper Paleolithic genome [11].

## Estimates for Europeans and East Asians

Our primary results (Fig 4) were obtained from eight diploid genomes of European and East Asian individuals (two each French, Sardinian, Han, and Dai) using our standard parameter settings (see above and Methods). For all real-data applications, to minimize noise from the randomized elements of the procedure (namely, coalescent simulation and generation of the perturbed calibration map), we averaged 25 independent calibrations of the data to obtain our final point estimate. With all eight individuals combined, we estimated a mutation rate of *μ* = 1.61 ± 0.13 × 10^−8^ per generation (Fig 4A). Using this value of *μ*, our starting heterozygosity *H_S_*(0) ≈ 7.4 × 10^−5^ corresponds to a TMRCA of approximately 1550–3100 generations, or 45–90 ky, assuming an average generation time of 29 years [12].

**Figure 4.**
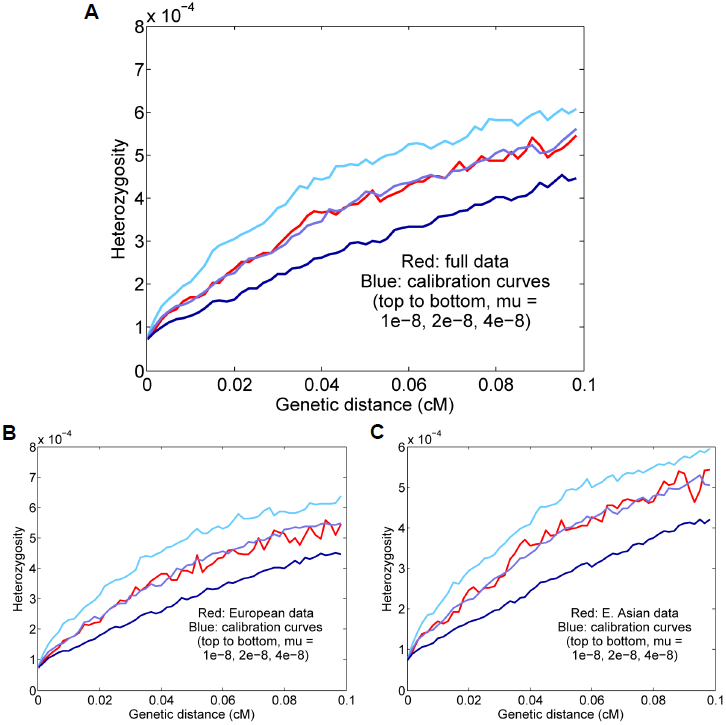
Results for Europeans and East Asians. (A) All eight individuals together; the inferred rate is *μ* = 1.61 ± 0.13 × 10^−8^ per generation. (B) Results for the four Europeans; the inferred rate is *μ* = 1.72 ± 0.14 × 10^−8^. (C) Results for the four East Asians; the inferred rate is *μ* = 1.55 ± 0.14 × 10^−8^. For all real-data results, the curves displayed are for representative calibrations matching the overall means. The reported values are also corrected for gene conversion, genotype error, and base content, which explains the apparent discrepancy between the final estimates and the curves (for example, the estimate (A) is corrected from a raw value of 2.00 × 10^−8^).

It is possible that our full estimate could be slightly inaccurate due to population-level differences in either the fine-scale genetic map or demographic history (see S1 Text). However, we expect Europeans and East Asians to be compatible in our procedure both because they are not too distantly related and because they have similar population size histories [19,24]. To test empirically the effects of combining the populations, we estimated rates for the four Europeans and four East Asians separately (Fig 4B–C). Using the same genotype error corrections, we found that the *H*_5−10_(*d*) curves as well as the final inferred values were similar to those for the full data: *μ* = 1.72 ± 0.14 × 10^−8^ for Europeans and *μ* = 1.55 ± 0.14 × 10^−8^ for East Asians. Thus, in conjunction with our simulation results, it appears that the full eight-genome estimate is robust to the effects of population heterogeneity.

Additionally, to investigate the influence of different mutational types, we estimated rates separately for CpG transitions and all other mutations (see Methods). We inferred values of *μ* = 0.50 ± 0.06 × 10^−8^ for CpGs and *μ* = 1.36 ± 0.13 × 10^−8^ for non-CpGs (S3 Figure), with a sum (1.87 ± 0.14 × 10^−8^) that is somewhat higher than our full-data estimate. Since CpG transitions are known to comprise approximately 17–18% of all mutations [8], our full-data and non-CpG estimates appear to be in very good agreement, whereas the CpG-only estimate is likely inflated, perhaps because our method performs poorly with the low density of heterozygous sites (only 1 per 100 kb window for our CpG-only starting points). As a result, we believe that our value of *μ* = 1.61 × 10^−8^ is accurate, or at most slightly underestimated, as a total mutation rate for all sites.

## Estimates for other populations

We also ran our procedure for three other non-African populations: aboriginal Australians, Karitiana (an indigenous group from Brazil), and Papua New Guineans. Using two genomes per population and computing curves for starting regions with 1–15 heterozygous sites per 100 kb (to increase the number of test regions, with a potential trade-off in accuracy), we inferred rates of *μ* = 1.86 ± 0.19 × 10^−8^, *μ* = 1.37 ± 0.19 × 10^−8^, and *μ* = 1.62 ± 0.17 × 10^−8^ for Australian, Karitiana, and Papuan, respectively (Fig 5). We note that the relatively high (but not statistically significantly different) per-generation value for Australians is consistent with the high average ages of fathers in many aboriginal Australian societies [12, 25]. Overall, given the expected small differences for historical, cultural, or biological reasons (including, as mentioned above, our use of the same “shared” genetic map for all groups), we do not see evidence of substantial errors or biases in our procedure when applied to diverse populations.

**Figure 5.**
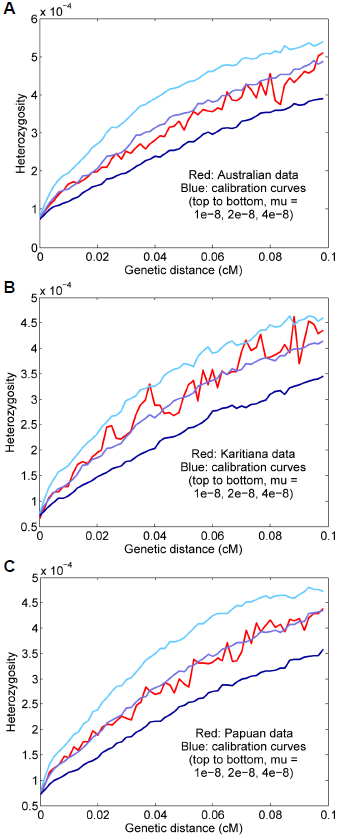
Results for other populations. (A) Australian, *μ* = 1.86 ± 0.19 × 10 ^−8^. (B) Karitiana, *μ* = 1.37 ± 0.19 × 10^−8^. (C) Papuan, *μ* = 1.62 ± 0.17 × 10^−8^.

## Discussion

Using a new method for estimating the human mutation rate, we have obtained a genome-wide estimate of *μ* = 1.61 ± 0.13 × 10^−8^ single-nucleotide mutations per generation. Our approach counts mutations that have arisen over many generations (a few thousand, i.e., several tens of thousands of years) and relies on our excellent knowledge of the human recombination rate to calibrate the length of the relevant time period.

We have shown that our estimate is robust to many possible confounding factors (S1 Fig). In addition to statistical noise in the data, our method directly accounts for ancestral gene conversion and for errors in genotype calls and in the genetic map. We have also demonstrated, based on simulations, that heterogeneity in demographic and genetic parameters, including the mutation rate itself, does not cause an appreciable bias. However, we acknowledge that our estimate requires a large number of modeling assumptions, and while we have attempted to justify each step of our procedure and to incorporate uncertainty at each stage into our final standard error, it is possible that we have not precisely captured the influence of every confounder. Similarly, while we consider a broad range of possible sources of error, we cannot guarantee that there might not be others that we have neglected.

### The meaning of an average rate

It is important to note that the mutation rate is not constant at all sites in the genome [26]. As we have discussed, we believe that this variability does not cause a substantial bias in our inferences, but to the extent that some bases mutate faster than others, a rate is only meaningful when associated with the set of sites for which it is estimated. For example, methylated cytosines at CpG positions accumulate point mutations roughly an order of magnitude faster than other bases because of spontaneous deamination [3,7,8,10]. Such effects can lead to larger-scale patterns, such as the higher mutability of exons as compared to the genome as a whole [27].

In our work, we filter the data substantially, removing more than a third of the sites in the genome. The filters tend to reduce the heterozygosity of the remaining portions [24,28], which is to be expected if they have the effect of preferentially removing false heterozygous sites. We also make a small adjustment to our final value of *μ* to account for differences in base composition between our ascertained starting points and the (filtered) genome as a whole (see Methods). For reference, in S1 Table, we give heterozygosity levels and human–chimpanzee divergence statistics for sites passing our filters, i.e., the subset of the genome for which our inferred rates are applicable.

### Evolutionary implications and comparison to previous estimates

A key application of the mutation rate is for determining the divergence times of human populations from each other and of humans from other species [1]. Published mutation rate estimates are highly discrepant, however, by as much as a factor of 2 between those based on sequence divergence among great apes (2−2.5 × 10^−8^ per base per generation) versus *de novo* mutations in present-day families (generally 1−1.2 × 10^−8^) [1–3]. This uncertainty causes estimated population split times to be highly dependent on whether a high or low rate is assumed. We also note that while the *de novo* mutation rate and the long-term substitution rate are equal under a neutral model (and assuming a constant value of *μ* over time), this may not hold in reality. In this case, one might expect rates measured over longer time scales to be somewhat lower, due to purifying selection, with our estimate affected more than *de novo* values but less than inter-species comparisons.

Here, we infer an intermediate rate of 1.61 ± 0.13 × 10^−8^ per base per generation, in good agreement with a previous estimate based on linked microsatellite mutations (1.4−2.3 × 10^−8^) [13] and a concurrent study using mutations present in inferred identical-by-descent segments (1.66 ± 0.04 × 10^−8^) [18]. Assuming an average generation time of 29 years [12] for the last ~ 50−100 ky (the time period over which our rate is measured), this value equates to 0.55 ± 0.05 × 10^−9^ per base per year (which overlaps the high end of the range of 0.4−0.6 × 10^−9^ inferred from comparisons between modern samples and a Paleolithic modern human genome [11]). Here we make the approximation that because most mutations are paternal in origin and accumulate roughly linearly with the age of the father [8, 10, 29], the per-year mutation rate is more robust to changes in the generation interval than is the per-generation rate. We stress that this conversion is approximate, both because the generation interval is uncertain and because this model is overly simplistic [3], but it allows us a reasonable means to compute long-term population split times even if the generation interval has changed in the past.

We propose that split times derived from our inferred per-year rate are in good agreement with the available paleontological evidence. Importantly, great apes exhibit a large amount of incomplete lineage sorting, which is indicative of large population sizes and high rates of polymorphism in their common ancestors [30,31]. This results in substantial differences between genetic divergence times and the final split times of species pairs. For example, according to a recent estimate [31], assuming a “fast” rate of 1.0 × 10^−9^ mutations per base per year (2.5 × 10^−8^ per base per generation with a generation interval of 25 years) results in an estimated population split time of ~ 3.8 million years (My) for humans and chimpanzees—with a genetic divergence time approximately 50% older— which seems too recent in light of the fossil record. By contrast, our rate of 0.55 × 10^−9^ leads to a more reasonable split time of 6.8 ± 0.6 My (or 7.3 ± 0.6 My using our filtered subset of the genome, with 1.23% sequence divergence; see above and S1 Table). Another constraint is the human–orangutan split, which is believed to be no older than about 16 My [1, 32]. In this case, our rate implies a split time of 20.1 ± 1.7 My (with a genetic divergence time substantially older, approximately 29 My) [31]. Although this appears to predate the fossil-inferred split (with some uncertainty), it is reasonable to expect that some changes in the biology (and, specifically, the mutation rate) of ancestral apes have occurred over this time scale (and likewise for older splits [1]). By comparison, however, a rate of ~ 0.4 × 10^−9^ per year from *de novo* studies implies a much more discrepant split time of ~ 28 My.

Our results can also be assessed in terms of their implications for split times among modern human populations. It has been argued on the basis of results from demographic models that a slower rate fits better with our knowledge of human history [1, 33]. For example, a recent method for estimating population split times from coalescent rates placed the median split of African from non-African populations at 60–80 ky and the split of Native Americans from East Asians at ~ 20 ky, both assuming a per-generation mutation rate of 1.25 × 10^−8^ and an average generation interval of 30 years [33]. While both the model and the histories of the populations involved are somewhat complicated, it does seem unlikely that these dates could be half as old (30–40 ky and 10 ky), as would be required for a rate of 2.5 × 10^−8^. Using our inferred rate also makes the dates more recent, but only modestly so: ~ 47−62 ky and 15 ky, with some associated uncertainty both from the model and from our estimated rate. Emphasizing the degree to which genetic variation is shared among modern humans, the genome-wide average heterozygosities of our samples (after filtering) range from 5.4−7.5 × 10^−4^, which corresponds to average TMRCAs of 480–670 ky between two haploid chromosomes (S1 Table).

A possible explanation for the discrepancy between our results and those of trio sequencing studies is that because it is very difficult to separate true *de novo* mutations from genotype errors in single-generation data, some mutations have been missed in previous work. For example, three recent exome-sequencing studies [34–36], which estimated effective genome-wide mutation rates of approximately 1.5 × 10^−8^, 1.2 × 10^−8^, and 1.35 × 10^−8^ per base per generation, found in follow-up validation that, in addition to virtually all sites from their filtered data sets, many putative sites that did not pass all filters (roughly 70%, 20%, and 90% of tested sites, respectively) were confirmed as true *de novo* mutations. These results suggest that there may be a subset of *de novo* mutations having low quality metrics that are missed in trio-based counts as a result of the filtering that is necessary to remove genotype errors. It would be of interest to carry out larger-scale follow-up validation to test if this is the case. In theory, filtered sites can be accounted for by adjusting the denominator in the final rate calculation [10], but it seems possible that site-level filters preferentially remove *de novo* mutations or that the resulting denominators have not been fully corrected, or both.

Another possibility is that the mutation rate could have changed over time. It has been suggested, for example, that phylogenetic and pedigree-based estimates could be reconciled if the rate has recently slowed in extant great apes [1]. While there is evidence of recent increases in the frequency of certain mutation types in Europeans [37], it seems unlikely that such changes have caused substantial differences in the total mutation rate over the last 50–100 ky. On the other hand, at least part of the discrepancy between our results and previous estimates could plausibly relate to environmental or cultural effects. For example, present-day hunter-gatherers, who may serve as the best available population proxy for our long-term rate estimate, have high paternal ages on average (about 32.3 years) [12]. Using an estimate from a recent *de novo* study [10] that each additional year of paternal age results in an average of 3.9 × 10^−10^ more mutations per base per generation, the sex-averaged mutation rate would be expected to be about 0.08 × 10^−8^ higher for a population with an average paternal age of 32.3 years than for the individuals in that study (average paternal age of 28.4 and inferred mutation rate of 1.27 ± 0.06 × 10^−8^ [10]), accounting for more than 20% of the discrepancy with our inferred value of 1.61 ± 0.13 × 10^−8^. One could also speculate that changes in lifestyle, diet, nutrition, or other environmental factors could potentially have contributed to a reduction in the mutation rate in the very recent past. In the future, we expect that new data, including from more diverse populations, combined with new technologies and analytical techniques, will continue to add to our knowledge of the human mutation rate, both in the precision of estimates of its long-term average and in its variability over time, by age and sex, and in different human groups.

## Methods

### Definition of the statistic *H_S_*(*d*)

Our inferences are based on a statistic that allows us to compare the mutation rate to the (much better measured) recombination rate. Intuitively, we compare local levels of heterozygosity in diploid genomes to the distance scales over which these levels change; the former are proportional to the mutation rate and the latter to the recombination rate, with the same constant of proportionality.

We define a statistic *H_S_*(*d*) equal to the average heterozygosity (local proportion of heterozygous sites) as a function of genetic distance *d* (measured in cM) from a set S of suitably ascertained starting points within a collection of diploid genomes (see below for more details). This statistic takes the form of a relaxation curve, with the rate of relaxation informative of the average TMRCA at the starting points (via the recombination rate), and the starting heterozygosity in turn informative of the mutation rate. In practice, we compute *H_S_*(*d*) in 60 distance bins from 0 to 0.1 cM.

### Locating starting points

In order to maximize signal quality, we would like to measure *H_S_*(*d*) using many starting points in the genome, but within a relatively narrow range of local heterozygosity at those points. For our primary analyses, we use a starting heterozygosity value *H_S_*(0) ≈ 7.5 × 10^−5^, corresponding to points with TMRCA roughly one-tenth of the genome-wide average for non-African individuals. This choice of S has two main advantages (see more detailed discussion below). First, there are relatively many such points in genomes of non-African individuals because it corresponds approximately to the age of the out-of-Africa bottleneck. Second, a smaller *H_S_*(0) corresponds to a slower and higher-amplitude (from a lower starting value to the same asymptote) relaxation of *H_S_*(*d*), making the curve easier to fit and less susceptible to genetic map error, an important consideration for our method (see below).

To determine local heterozygosity levels, we tile the genome with 100-kb windows (defined by physical position) and count the proportion of heterozygous sites within each (after filtering, such that that each window will have fewer than 100, 000 un-masked sites). The starting points used to compute *H_S_*(*d*) are then the midpoints of the 100-kb regions having a heterozygosity at the desired level, for example 5−10 × 10^−5^ for out-of-Africa-age blocks with *H_S_*(0) ≈ 7.5 × 10^−5^. We use the notation (for example) H_5−10_(d) to denote an *H_S_*(*d*) curve computed for starting points with 5–10 heterozygous sites per 100 kb. This scheme may result in choosing starting points with unwanted true heterozygosity if there are recombinations within the 100-kb region, but 100 kb is long enough that most regions within a narrow range of heterozygosity on that scale should be similarly behaved. Additionally, any deviations from the desired heterozygosity range at the midpoints should, on average, be the same for real and simulated data (assuming that the simulator accurately models the genealogical process with recombination; see below), and hence would only cause noise rather than bias in the estimated mutation rate. Similarly, while randomness in the number of accumulated mutations causes the relationship between observed heterozygosity and TMRCA to deviate from strict linearity, this effect is the same in real and simulated data. As an attempt to avoid certain kinds of undesirable behavior (for example, a very low heterozygosity over most of the region and a recombination near one end followed by high heterozygosity), we also require at least one heterozygous site in each half of the window (except for the CpG-only results; see below). Descriptive statistics for regions ascertained with *S* = 5 − 10 can be found in S2 Table.

### Inference strategy

As described above, *H_S_*(*d*) exhibits a relaxation as a function of *d* as a result of ancestral recombination events. Recombination can be modeled as a Poisson process (in units of genetic distance), but *H_S_*(*d*) does not have an exponential functional form, because the TMRCAs *T*_1_ and *T*_2_ at two loci separated by a recombination event are not independent [19, 38, 39]. First, both *T*_1_ and *T*_2_ must be older than the time at which the recombination occurred, which imposes different constraints on *T*_2_ for different values of *T*_1_ This dependence becomes especially complicated when the population from which the chromosomes are drawn has changed in size over time. Second, the coalescence at time *T*_2_ can involve additional lineages in the ancestral recombination graph, making the expected time different than would be true for two lineages in isolation. For example, with some probability, the two lineages split by the recombination can coalesce together more recently than this combined lineage coalesces with the second chromosome, in which case *T* = *T*_2_.

These complicating factors mean that *H_S_*(*d*) is difficult to describe as a closed-form function of *d*. However, we know that *H_S_*(*d*) relaxes from *H_S_*(0) toward the average heterozygosity 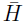, and the rate of relaxation is governed by the relationship between *μ* and *r*. Thus, our strategy is to infer the true value of *μ* by simulating sequence data matching our real data in all respects (see below for more detail) and with a range of different values of *μ* (typically *μ* = 1, 2, 4 × 10^−8^). Then, we can compare the observed *H_S_*(*d*) curve to the same statistic calculated on each simulated data set and infer *μ* by finding which value gives the best match. Computationally, we interpolate the observed *H_S_*(*d*) curve between the simulated ones (from *d* = 0 to 0.1 cM), parametrized by the simulated *μ* values (we use the MATLAB “spline” interpolation; for our main results, other interpolation methods yield results differing by less than 1%). Finally, we perform variance-weighted least-squares to find the single best-fit value of *μ*.

### Population size history

We estimate the historical population sizes for the sampled chromosomes with PSMC [19]. PSMC returns parameters in coalescent units: the scaled mutation rate *θ* = 4*Nμ*, the scaled recombination rate *θ* = 4*Nμ*, and population sizes going back in (discretized) time, with both the sizes and time intervals in terms of the scaling factor *N* (the baseline total population size). We do not know *N*, but the inferred *θ* together with the population size history are exactly what we need in order to simulate matching data for the calibration curves. We do not use the inferred value of *ρ* but rather set *ρ* = *θ*r/*μ*, where *r* is the true per-base recombination rate and *μ* is the fixed mutation rate for a given calibration curve. This maintains the proper ratio between *r* and *μ* for that curve, as well as the proper diversity parameter *θ*. While we only use short regions of the genome in computing *H_S_*(*d*), we run PSMC on the full genome sequences. The exception is that when testing the method in simulations, we run PSMC on the simulated segments, as these are all that are available to us (see below). Further technical details can be found in S1 Text.

### Genetic map error

The statistic *H_S_*(*d*) is computed as a function of genetic distance, which we obtain from a previously-estimated genetic map. However, while map distances (i.e., local recombination rates) are known much more precisely than mutation rates, there is still some error in even the best maps, which we must account for in our inferences.

Our approach is not to make a direct correction for map error but rather to include it in a matching fashion in the calibration data. We first select a baseline genetic map from the literature, and we plot *H_S_*(*d*) as a function of *d* using this base map as the independent variable. To make the calibration curves match the real data, whose intrinsic, true map does not match the base map exactly, we simulate the calibration data using a perturbed version of the base map, with the aim of capturing an equal amount of deviation as in the real data.

The base map we use is the “shared” version of an African-American (AA) genetic map published in [17]. The AA map was derived by tabulating switch points between local African and European ancestry in the genomes of African-Americans, which reflect recombination events since the time of admixture. The “shared” component of the map was estimated as the component of this recombination landscape that is active in non-Africans, particularly Europeans. We also considered using either the 2010 deCODE map [16], which was estimated by observing crossovers in a large Icelandic pedigree cohort, or an LD map estimated from variation in linkage disequilibrium levels in unrelated individuals [14, 15]. However, we were concerned that an LD-based map would cause confounding in our method because it is based on a correlated ancestral recombination signal, while we found that at short distance scales, the “shared” map was less susceptible than the deCODE map to the issue of zero-length intervals (see prior distribution below).

While using as accurate a map as possible is helpful, the key to our approach is in the ability to quantify the amount of error in the base map. In what follows, we describe in detail our methods to measure the degree of genetic map error.

#### Basic model and previous estimates

Our basic error model is that of [20]. Consider a chromosomal interval whose true genetic length is *Z*. Given the measured genetic length *g* of the interval in our base map, we assume that *Z* ~ Gamma(*αg*, *α*), so *E*[*Z*|*g*] = *g* and var(*Z*|*g*) = *g*/*α*. The gamma distribution has several desirable properties, in particular that it is scale-invariant. The parameter *α* measures the per-distance variance of the map (and hence of our perturbation), with *α* larger value of α corresponding to a smaller variance. It can be interpreted directly as the inverse of the length scale at which the coefficient of variation of the map equals 1: on average, intervals of length 1/*α* have an equal mean and standard deviation, while longer intervals are relatively more accurately measured and shorter intervals are less accurate.

Briefly, this model can be used to estimate *α* by comparing genetic distances in the base map to counts of recombination events observed in an independent validation data set: the more accurate the map, the more closely it will predict the observations. From the distribution above, one can specify a full probability model for the validation data in terms of the error in the base map as well as other parameters and infer *α* via a Bayesian (Gibbs sampling) approach. Full details can be found in [20]. In that study, using a validation data set of recombination events observed in a Hutterite pedigree [40], the authors obtained estimates (in per-Morgan units) of *α* = 1400 ± 100 M^−1^ for the deCODE map [16] and *α* = 1220 ± 80 M^−1^ for the HapMap LD map [15].

Here, we apply the same method to estimate *α*, but using a much larger set of validation data, consisting of 2.2 million crossovers from 71, 000 meioses in Icelandic individuals [23] (versus 24,000 crossovers from 728 meioses in [40]). A potential complication is that the procedure to build the “shared” map [17] used information from the 2010 deCODE map, which is not independent from our validation set. However, we reasoned that we could constrain the true α by applying the estimation method first to the “shared” map as an upper bound (since the value will be inflated by the non-independence) and then to the full AA map as a lower bound (since the AA map includes African-specific hotspots not active in Europeans and does not use information from the deCODE map).

As noted above, since the “shared” map is estimated from directly-observed crossover events rather than population-level statistics (as for LD-based maps), it should be free from confounding between heterozygosity and recombination (as desired for our rate estimation method). There could potentially be a subtle effect of population heterozygosity on the SNP grid on which the map is defined, which does not have perfectly uniform coverage of the genome, but both the base map data and validation data should be independent. Moreover, the error model described above takes into account differential SNP spacing [20]. Also, we note that most of the power of our procedure comes from regions away from recombination hotspots, so that any potential issues pertaining to hotspots, including differential SNP density, should be minimized.

#### Modified prior distribution on *Z*

We also add one modification to the basic model of map error described above. For intervals in the base map with estimated (genetic) length 0, the original model states that the true length of these intervals must be 0 (since *Z* has mean *g* = 0 and *Z* ≥ 0). In fact, though, the data used to build the map might simply have included no crossovers there by chance. Overall, very short intervals in the base map are likely underestimated, while very long intervals are likely overestimated.

To account for this effect, we modify the prior distribution on the true length *Z* by adding a pseudo-count adjustment *π*, i.e., a small uniform prior on the true map length. In order for the model still to be additive, it is reasonable for the prior to be in units of genetic distance per physical distance.

Empirically, we observe that without the adjusted prior, the relaxation of *H_S_*(*d*) in the calibration curves is too slow at the smallest values of *d* and too fast at larger *d* (S4 Fig). This would be expected if very short intervals in the base genetic map are underestimated, so that the calibration data have too few recombinations in that range compared to real data. By matching the curve shape of real data to the calibration data for different values of the pseudo-count, we find that a value of *π* = 0.09 cM/Mb properly corrects for the underestimation of very short intervals in the “shared” map.

#### Perturbed genetic maps for calibration data

For our purposes, once we obtain a value for *α*, we use the gamma-distribution model to generate the randomized perturbed maps that we input to msHOT to generate the calibration data (see below). Our complete error model is as follows: for an interval of physical length *p*, we take *Z* ~ Gamma(*αg*′, *α*), where *g*′ = *γ*(*g* + *πp*) for a pseudo-count n and the corresponding constant factor *γ* < 1 that preserves the total map length. Thus, for each interval in the map (between two adjacent SNPs on which the map is defined), if the physical length is *p* and the genetic length in the base map is g, we create a new genetic length *Z* for that interval in the perturbed map by drawing from this distribution. When computing *H_S_*(*d*), we then measure all genetic distances in both the real and simulated data by linearly interpolating individual sites within the SNP grid.

#### Uncertainty in *α*

The importance of accurately estimating α lies in the fact that the smaller the value of *α* used to generate the calibration data (i.e., the less accurate the genetic map is taken to be), the smaller the final inferred value of *μ* will be. This is because the genetic lengths used for simulation will be more discrepant from the base map, and as a result, the final value of *H_S_*(*d*) (computed on the calibration data) for a given *d* will reflect an average of the heterozygosity over a wider range of perturbed distances around *d*. Since *H_S_*(*d*) is a concave function of *d*, this smoothing will cause *H_S_*(*d*) to decrease, or in other words, to make the relaxation appear slower. Thus, in order to match the real data, a calibration curve with a smaller *α* would have to have a higher scaled recombination rate, and hence, with a fixed *θ*, a smaller *μ*.

As described in Results, we assume a standard error of 300 around our estimate of *α* = 3100 M^−1^. To determine how much of an effect this uncertainty has on our estimates of *μ*, we run our procedure with our primary data set using a range of different values of *α* (2500, 3000, 3500, 4000, and 4500; see S2 Fig). The slope of a linear regression of the inferred *μ* as a function of *α* is 1.66 × 10^−4^ (in units of 10^−8^ per base per generation per M^−1^), so that the standard error of 300 for *α* translates into a standard error of roughly 0.05 × 10^−8^ per base per generation for *μ*.

We also note that even if our map error model is properly specified and estimated, there could be a small bias in our final inferred *μ* if the exact form of the true map is such that the real-data *H_S_*(*d*) curve relaxes slightly faster or slower than the calibration curves built with a perturbed map. Since this uncertainty is analogous to variability in the inferred *μ* depending on the exact instantiation of the perturbed map, we can compensate for it by using for our final point estimate the average of calibration results for different versions of the perturbed map. In practice, we obtain our final point estimates by averaging 25 independent calibrations of the data, which should remove most randomization noise arising from both the perturbed map and the simulations, and we assume that the resulting reduction in uncertainty (as compared to our single-run jackknife standard error) negates the uncertainty from the exact form of the true map.

### Simulation of calibration data with msHOT

As discussed above, the relaxation of *H_S_*(*d*) reflects the decorrelation of heterozygosity (caused by recombination) as a function of genetic distance. However, in the sequence of TMRCAs for the recombination-separated blocks along a chromosome, successive values are not independent, and in fact the sequence is not Markovian, since even lineages that are widely separated along the chromosome can interact within the ancestral recombination graph [38, 39]. It is important, then, that our simulated data be generated according to an algorithm that captures all of the coalescent details that could impact the history of a real-data sample. For this reason, we use msHOT [41] (an extension of ms [42] that allows variable recombination rates) rather than a Markovian simulator, which would have had the advantage of greater speed.

As a result of using a non-Markovian simulator, it is computationally infeasible to generate entire simulated chromosomes. Thus, in practice we define wider “super-regions” around the 100-kb windows and simulate the super-regions independently of each other, matching the physical and genetic coordinates to the human genome. Since we compute *H_S_*(*d*) from *d* = 0 to 0.1 cM, we define the super-regions to include at least 0.1 cM on both sides of their internal starting point, which typically leads to a total length of several hundred kb per super-region.

Finally, in addition to matching the demographic and genetic map parameters of the calibration data to the test data, we also apply an adjustment to the calibration curves themselves to correct for residual unequal heterozygosity (not precisely captured by PSMC), which would cause the asymptotes of the curves to be mis-aligned. In particular, we multiply the relaxation portion of the calibration curves (i.e., *H_S_*(*d*) – *H_S_*(0)) by the ratio of the heterozygosity of the real data (over all of the super-regions) to that of the matching simulated data. In our experience, this correction ranges from 0–10%. We note that in theory, the intercept values *H_S_*(0) might also not be recapitulated exactly in the calibration data, but in fact the intercepts match extremely closely in all cases other than the CpG-only estimate (see below). This indicates good reconstruction of the demographic history around the TMRCAs of our test regions.

### Genotype error

When we compare matching *H_S_*(*d*) curves for real and simulated data, the real-data curve will be influenced by genotype errors (almost all of which are sites that are in fact homozygous but are mistakenly called as heterozygous). Since the relaxation rates of the curves as a function of *d* are equal, the starting points will have the same average TMRCAs. The real and simulated regions also have the same average starting heterozygosity *H_S_*(0), but since the real data contain both true heterozygous sites and genotype errors, the calibration data must have a higher mutation rate per generation. Thus, false-positive heterozygous sites will artificially inflate our estimates of *μ*. The upward bias in the estimates will be larger for smaller values of *H_S_*(0), since the local ratio of false to true heterozygous sites will be larger; for the same reason, for a fixed error rate, our method will be less sensitive to genotype errors than *de novo* mutation counts.

We have two main approaches for dealing with errors in genotype calls. First, we have taken a number of steps to filter the data, discussed below, such that the sites we analyze have high-quality calls and are as free from errors as possible. However, this is not sufficient to eliminate all false positives, and thus we also use local abundances of CpG transitions to quantify the proportion of true homozygous sites that are called as heterozygous (for full details, see S1 Text). For our final values of *μ*, we directly correct for these inferred levels of genotype error, incorporating our uncertainty in the error rate into our final reported standard error. Specifically, for a given starting heterozygosity value *H_S_*(0) and genotype error rate *ε* (inferred for the same set of windows S), we multiply the initial estimate of *μ* by a factor of (*H_S_*(0) – *ε*)/*H_S_*(0). This correction is based on the idea stated above that while the calibration data are created to mimic the full observed data (including genotype errors), matched curves with the same *H_S_*(0) and slope of relaxation (and thus TMRCA) will still differ: the real data will have errors contributing to the local heterozygosity, so that the inferred mutation rate from the calibration will be too high. (The same logic applies for the two sections that follow; see also S1 Text). To ensure that this correction is valid, we perform simulations in which we randomly add false heterozygous sites to the simulated sequence data (see below).

### Non-crossover gene conversion

Gene conversion is similar to recombination (as we have used the term; more precisely, crossing-over) in that it results from double-stranded breaks during meiosis and leads to the merging of genetic material between homologous chromosomes. However, whereas crossing-over creates large-scale blocks inherited from the recombining chromosomes, noncrossover gene conversion occurs in very small tracts, on the order of 100 bases in humans [43]. For our purposes, gene conversion is significant primarily because it can introduce heterozygous sites in our test regions if one of the haplotypes has experienced a gene conversion event since the TMRCA. (New mutations that occur in our regions can also be lost via gene conversion, which we take into account, although this rate is much lower because the local heterozygosity is small.)

We choose to account for the effect of gene conversion by applying a correction to our inferred mutation rates, reasoning that a subset of the observed heterozygous sites will be caused by gene conversion events rather than mutations. Since our method relies on the ratio between the recombination and mutation rates at the selected starting points, the key quantity is the number of heterozygous sites near those points that are due to gene conversion, which we subtract from the raw estimate of *μ* after correcting for genotype error. The gene conversion rate is a combination of two factors: the probability that each base is involved in a gene conversion event and the conditional probability that a polymorphism is introduced. For the former, we use a recent estimate that non-crossover gene conversion affects approximately 5.9 × 10^−6^ bases per generation (95% confidence interval 4.6−7.4 × 10^−6^) [21] and adjust for local recombination rate, and for the latter, we use differences between the heterozygosity in the test regions and in the genomes as a whole (see S1 Text). The final correction ranges from 0.13−0.17 × 10^−8^ per base per generation (with a standard error of 0.02−0.03 × 10^−8^), approximately 7–10% of the total apparent mutational signal after accounting for genotype error. To confirm that this procedure is accurate, we also apply it to simulated data with gene conversion (see below and S1 Text).

We note that gene conversion events are well known to carry a GC bias, so that when they affect heterozygous sites with one strong (C or G) and one weak (A or T) allele, the strong base is preferentially transmitted (recently estimated as 68±5% of the time [21]). In theory this bias could have an impact on our gene conversion correction, but (a) genome-wide, weak-to-strong and strong-to-weak SNPs are close to equally frequent (the latter about 10% more abundant), and (b) while new mutations are enriched in strong-to-weak substitutions, gene conversion of newly occurring mutations is rare in our test regions because of their recent TMRCAs. Thus, for simplicity, other than a small adjustment when calculating the gene conversion corrections for the CpG-only and non-CpG-only estimates, we do not include GC bias in the gene conversion correction (we note that this is conservative, in the sense that GC bias would cause the gene conversion effect to be slightly weaker than we are assuming).

### Base content mutability adjustment

Because not all bases in the genome are equally mutable, and our test regions may deviate from the full genome in their base composition, we apply an adjustment to our estimated rates to convert them to a genome-wide equivalent. The two primary parameters we use are the fraction of CpG sites (which are highly mutable) and total GC content (because *g* and C are more mutable than A and T, beyond CpG effects). We also include an interaction term between these two quantities. Our final adjustment for the primary eight-genome estimate is a multiplicative factor of 1.027 ± 0.003 (an increase of about 0.04 × 10^−8^ over the uncorrected value). The uncertainty in the adjustment incorporates confidence intervals on the parameters in our model and a jackknife for the counts of CpG and GC sites. Full details can be found in S1 Text.

### Choice of the starting heterozygosity range S

Our use of *S* = 5−10 per 100 kb for our primary results was guided by several considerations. First, our method works best with relatively recently coalesced starting points (at most perhaps 1/3 of the genome-wide average) because they yield *H_S_*(*d*) curves that relax slowly and have a low intercept *H_S_*(0). At a finer scale, we also wish to to balance the impact of different sources of error: while genetic map error is more significant for higher starting heterozygosity, genotype error is relatively more significant for lower starting heterozygosity. Another consideration is that the TMRCAs corresponding to *S* = 5−10 lie approximately at the deepest portion of the out-of-Africa bottleneck [19, 33], so that more starting points are available at that range (for non-African populations). Moreover, demographic inference with PSMC is less accurate both at very recent times and at the edges of bottlenecks [19]. The combination of these factors motivates our use of *S* = 5−10.

### Noise and uncertainty

Several of the steps in our procedure have some associated statistical uncertainty, while others rely on randomization. These include the computation of *H_S_*(*d*) from a finite number of loci, the population size inference with PSMC, the random genetic map perturbation, and the simulation of calibration data. In order to capture this uncertainty, we use jackknife resampling to obtain standard errors for our estimates of *μ*, treating each autosome as a separate observation and leaving out one chromosome in each replicate. Our rationale for this scheme is that nearby regions of a chromosome are non-independent, and different individuals can also have correlated coalescent histories for a given locus, but a chromosomal unit encompasses most or all of the dependencies among the data. For real data, we also average the results of 25 independent calibrations to obtain our point estimate of *μ*, which eliminates most of the noise associated with randomization. We note that we have found that the transformation from PSMC-inferred demographic parameters to calibration data via msHOT is discontinuous and is not properly captured by the jackknife. Thus, we use the same population size history (from the full data) for each replicate (see next section and Results).

The other form of uncertainty in our method is our inexact knowledge of a number of model parameters, including the genetic map variance α and several components of the three final adjustments (genotype error, gene conversion, and base content; see S1 Fig). For these sources of error, we translate our uncertainty in the parameters into uncertainty in *μ* and combine them with our jackknife results to form a final standard error, using the assumption that the errors are independent and normally distributed (for implicit conversion of standard errors into confidence intervals).

### Simulations

To test the accuracy of our procedure in a controlled setting, we first apply it to simulated data generated using msHOT [41]. When not otherwise specified, we create 20 sample genomes with *μ* = 2.5 × 10^−8^; an ancestral population size of 10,000 (or 16,666 when using *μ* = 1.5 × 10^−8^ so as to maintain the same diversity parameter *θ*) outside of a 10 × bottleneck from 1000–2000 generations ago (similar to the age of the out-of-Africa bottleneck); and a perturbed version of the “shared” AA genetic map (*α* = 3000 M^−1^ and n = 0.09 cM/Mb). We run the full inference procedure as we would with real data, except with a default total of 30 genomes’ worth of data per calibration curve versus 40 for real data. Also, for computational efficiency, when running PSMC on simulated test data, we only include a single copy of each chromosome (chosen at random from among the samples in the simulated data set).

As discussed in more detail in S1 Text, in addition to this basic setup (a), we also run a number of additional simulations. First, we run the procedure (b) with a true rate of *μ* = 1.5 × 10^−8^, (c) with a true rate of *μ* = 1.5 × 10^−8^ plus gene conversion, and (d) with simulated genotype errors. Then, to test the effects of possible model violations, we simulate (e) samples from an admixed population, (f) mutation rate heterogeneity based on polymorphism levels in present-day African individuals, and (g) all three complications (d)-(f) simultaneously. For simulations (d) and (g), we add false heterozygous sites to the simulated diploid genomes (at a rate of 1 per 100 kb for (d) and 1 per 150 kb for (g)) and apply our standard correction, with one modification: because msHOT does not create individual nucleotides, we directly count the numbers of errors in ascertained regions instead of using our CpG-based estimate.

### Data and filtering

As mentioned previously, we generate our estimates using genome sequences from non-African individuals, since the presence of a large number of relatively recently coalesced blocks arising from the out-of-Africa bottleneck gives us more data to work with at starting points with low heterozygosity. We use high-coverage sequences published in [24] and [28].

In order to remove as many genotype errors as possible, we use a filtering scheme based on the one applied to estimate heterozygosity in [28]. This consists of a tandem repeat filter, mapping quality threshold (MQ = 30), genome alignability filter (all possible 35-mers overlapping a given base match uniquely to that position in the genome, with up to one mismatch), and coverage thresholds (central 95% of the depth distribution) [28]. We additionally apply a strict genotype quality threshold in order to preserve the highest-quality calls for analysis. From the GATK output, we compare the PL likelihood score of the heterozygous state to the minimum of the two PL scores of the homozygous states, imposing a quality threshold of 60 along with a prior of 31 (to reflect the genome-wide average heterozygosity). That is, if the heterozygote PL is at least 60 + 31 = 91 lower than either homozygote PL, we call the site heterozygous; if it is at least 60 − 31 = 29 higher, we call the site homozygous; and if it is in between, we mask the site as low-quality. Finally, we also remove all sites 1 or 2 bases away from any masked base under the five filters described.

While filtering is not necessary for the simulated calibration data, we still apply the same filters to the calibration data as to the real sequence data for consistency, on a genome-matching basis (e.g., for a sample of eight real genomes and our default real-data setting of 40 genomes’ worth of calibration data, the base positions that are masked for each real sequence are also masked in five of the simulated sequences). In addition to filtering out individual sites, we impose a missing-data threshold for regions, ignoring any with more than 50% of sites masked (either of the super-region or the 100-kb central window).

### Estimates for separate mutation classes

For our estimates of the mutation rate for CpG and non-CpG sites separately, we make several small modifications to our default procedure. First, we divide heterozygous sites into two classes: C-to-T transitions at CpG sites (defined based on the human reference sequence) and all others. For the two estimates, we consider as homozygous all sites in the genome not in the corresponding class of heterozygous sites. This places the CpG, non-CpG, and full-data values on an equal denominator. For the CpG estimate, we select starting points with *S* = 0.1−2, which, because of our filtering, is equivalent to requiring exactly one heterozygous site in a 100 kb window. We then create calibration curves with *μ* = 0.2, 0.4, 0.8 × 10^−8^ and 10 simulated genomes per real-data sequence (two times the usual). We find that, unlike the curves for all of our other results, the CpG-only *H_S_*(*d*) has a noticeably lower y-intercept than the calibration curves, which we believe may be due to relatively poor reconstruction of the demographic history with thin data. In order to make the interpolation more sensible, we translate the calibration curves downward to match the real-data value of *H_S_*(0), which results in a decrease in the final inferred *μ* of approximately 0.04 × 10^−8^ as compared to the uncorrected fitting. For the non-CpG estimate, we use *S* = 4.375−8.75 and calibrate as for the full data.

When correcting for genotype error, we use our baseline value of 1.08 × 10^−5^ per base, multiplied by the fraction of CpG or non-CpG sites. Likewise, we partition the full-data gene conversion estimate by the fraction of heterozygous sites in each class and adjust for GC bias (final correction approximately 0.09 and 0.91 times the full gene conversion rate for CpG and non-CpG, respectively, versus about 0.12 and 0.88 for fractions of heterozygous sites). Finally, we re-compute base content corrections of approximately 1.17 for CpGs and 1.005 for non-CpGs (where the CpG estimate is more strongly affected because of the relatively low proportion of CpG sites in the ascertained regions).

### Software

`MATLAB` code is available at https://github.com/DReichLab/MutationRateCode.

## Acknowledgments

We thank Shop Mallick and Heng Li for their assistance with technical details and Guy Amster, Priya Moorjani, George Tucker, Pier Palamara, Shai Carmi, and Molly Prze-worski for helpful discussions. M.L. acknowledges support from the Simons Foundation and NIH (grant R01GM108348, to B.B.). P.L. was supported by NIH fellowship F32 HG007805 and S.S. by NIH grant K99 GM111744. N.P. and D.R. are grateful for support from NSF HOMINID grant #1032255 and NIH grant GM100233. D.R. is an Investigator at the Howard Hughes Medical Institute.

## S1 Text: Supplementary Methods

### Technical details for population size history inference

PSMC runs on a reduced version of the genome, with consecutive sites grouped into bins of 100 and each bin marked as 1 or 0 depending on whether there is at least one heterozygous site in the bin or not. Bins can also be marked as “missing” if a certain number of the 100 sites have un-called genotypes (90 in the original PSMC publication). We find that two aspects of this procedure can affect the overall average heterozygosity of simulated data generated from a PSMC-estimated population size history. First, different values of the missing-bin threshold lead to different heterozygosity levels. Second, the PSMC program accepts a maximum of one heterozygous site per bin, whereas there can in fact be more than one. This effect is non-linear as a function of TMRCA; since heterozygosity varies substantially along the genome, the program will systematically underestimate the age of the most anciently coalesced regions.

To account for these factors, we first use an empirically-determined threshold of 35 un-called sites per 100 to call a bin as missing, which yields a more closely matching final heterozygosity. Second, we implement a multiple-het-per-bin adjustment, whereby we modify the PSMC output before using it to create the calibration data, as follows. The population size history inferred from PSMC consists of discretized time intervals [*t t_i_*_+1_] (in coalescent units), with the population size *S_i_* in each interval telling us how much of the genome falls within that level of TMRCA (and hence of per-bin heterozygosity). Given these heterozygosity levels, we use a binomial distribution to scale the times defining the endpoints from the output units of probability of at least one heterozygous site per 100 bp to the implied expected number of heterozygous sites per 100 bp. Creating calibration data according to these new values more accurately recapitulates the true distribution of heterozygosity levels across the genome, as well as the total genome-wide heterozygosity.

Explicitly, if *θ* is the original baseline PSMC-inferred ms diversity parameter (per base), then we define the adjusted times *T_i_* (with T*_0_* = 0) and sizes *S_i_* recursively as

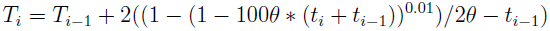
and 

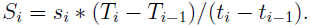

As an example, for our eight-genome PSMC inference, the population size trajectory (in ms notation) is adjusted from

> –eN 0.0049 0.1053 –eN 0.0080 0.0562 –eN 0.0118 0.0653 –eN 0.0162 0.1052 –eN 0.0214 0.1860 –eN 0.0277 0.3115 –eN 0.0350 0.4524 –eN 0.0438 0.5549 –eN 0.0542 0.5847 –eN 0.0664 0.5498 –eN 0.0810 0.4846 –eN 0.0983 0.4185 –eN 0.1188 0.3646 –eN 0.1431 0.3257 –eN 0.1718 0.3006 –eN 0.2060 0.2882 –eN 0.2464 0.2880 –eN 0.2944 0.3003 –eN 0.3513 0.3275 –eN 0.4187 0.3726 –eN 0.4987 0.4363 –eN 0.5935 0.5102 –eN 0.7059 0.5755 –eN 0.8391 0.6142 –eN 0.9971 0.6254 –eN 1.1844 0.6360

to

> –eN 0.0049 0.1053 –eN 0.0080 0.0564 –eN 0.0118 0.0655 –eN 0.0163 0.1060 –eN 0.0215 0.1874 –eN 0.0279 0.3152 –eN 0.0353 0.4587 –eN 0.0442 0.5653 –eN 0.0549 0.5978 –eN 0.0674 0.5655 –eN 0.0825 0.5012 –eN 0.1006 0.4364 –eN 0.1221 0.3834 –eN 0.1480 0.3464 –eN 0.1789 0.3237 –eN 0.2163 0.3153 –eN 0.2613 0.3210 –eN 0.3161 0.3424 –eN 0.3827 0.3839 –eN 0.4645 0.4517 –eN 0.5655 0.5513 –eN 0.6917 0.6788 –eN 0.8512 0.8168 –eN 1.0565 0.9467 –eN 1.3276 1.0732 –eN 1.6989 1.2608

As can be seen, the correction is negligible for more recent times where the density of heterozygous sites is low (and where our test regions have their TMRCA) and only becomes substantial for the oldest time intervals.

### Windows of measurement for correction factors

As discussed in Methods (“Genotype error”), because our method works by calibrating the mutation rate against the recombination rate via the local heterozygosity and non-recombined tract length at our starting points (equivalently, the intercept and slope of *H_S_*(*d*)), this local behavior is what we are concerned with when correcting for genotype error, gene conversion, and base content. These processes affect the entire *H_S_*(*d*) curve, but the calibration data recapitulate the full patterns of heterozygosity in the real data through the PSMC inference. Thus, once we fit the real-data curve to the calibration curves, which are ascertained to have similar *H_S_*(0), the question becomes how much this value *H_S_*(0) is impacted by various processes. While we would ideally like to derive the correction parameters directly at the starting points, in practice, we need to define windows around these points in order to obtain stable counts (as in the case of determining local heterozygosity in the initial ascertainment step). Aiming for a balance between capturing the behavior at the midpoint and maintaining high enough statistical power, we use windows of 10 kb when measuring the recombination rate for the gene conversion correction, 30 kb when measuring GC content and CpG site fraction for the base content mutability adjustment, and 50 kb when measuring CpG transition proportion for the genotype error correction (see below for more details for all three).

### Details of genotype error rate estimation

To estimate the genotype error rate, we take advantage of the fact that C-to-T transitions at CpG dinucleotides are greatly enriched among true mutations as compared to genotype errors. We assume that genotype errors are equally likely at CpG and non-CpG sites; while this may not strictly be true, the frequency of CpG sites in the genome is low enough (less than 2%) that deviations should only have a minor impact. More generally, we assume that the *a priori* probability of observing a genotype error is uniform across the genome. Because errors are rare, we do not have power to detect local variation in their rate of occurrence, and instead we aggregate over regions to produce as accurate an average error rate as possible.

For the fraction of CpG transitions among true heterozygous sites, we use a previous estimate of approximately 17.3% based on observation of almost 5000 de novo mutations [8]. Since GC-biased gene conversion results in a decrease in this frequency among older variants [3, 44], we use our gene conversion adjustment from above (roughly 0.14 × 10^−8^ per base per generation, with an order of magnitude more mutations introduced than removed for haplotypes in our TMRCA range), together with an estimated 2:1 transmission bias for GC bases [21], to estimate a reduction of approximately 0.6% for CpG transition frequency for our test regions. Given that our test regions have TMRCA roughly 10% of genome-wide average, this value is in line with previous reports of the difference in CpG transition fraction among *de novo* mutations versus common polymorphisms [3, 44]. To account for uncertainty, we use a standard error of 0.1% on the gene conversion correction (see below).

Using the same eight genomes as for our final *μ* estimates, with the same filtering and ascertainment of starting points, we compile the fraction of CpG transitions out of all heterozygous sites in the regions with local heterozygosity rates of 5–10 per 100 kb, with one caveat. Because CpG fraction strongly depends on the proportion of CpG sites, and we find that our test regions have relatively low such proportions (where we define CpG sites based on the human reference sequence), we truncate the low end of this distribution and only use test regions with at least 1.24% CpG sites (corresponding to 3551, or 37%, of the 9626 test regions), such that the overall proportion of CpG sites equals 1.85%, the same as in [8]. Additionally, we only use the central half of the 100 kb regions (i.e., 25 kb on either side of the starting points) to represent more precisely the local error rate at the starting points (while maintaining adequate statistical power). We then compare the observed CpG fraction (about 14.4%) to the gene conversion-adjusted expectation for true mutations (about 16.7%), from which we infer the presence of genotype errors at a rate of 1.08 × 10^−5^ per base (roughly 1 per 100 kb, or one-seventh of the heterozygous sites in our test regions). We note that the GC content in the thresholded regions is approximately 42% on average, which would imply a slight decrease in CpG mutability as compared to the genome-wide average (see base content adjustment below), but to be conservative (i.e., err on the side of slightly over-correcting), we do not make this adjustment here.

As described in Methods, we apply our correction by multiplying the raw inferred *μ* values by (*H_S_*(0) – *ε*)/*H_S_*(0), where e is the error rate. We incorporate this factor separately for each jackknife replicate, but using the same value of e throughout, as we found that re-computing the error rate within the jackknife was unstable. Instead, we account for our noisy estimate of the error rate by translating our uncertainty in e into uncertainty in *μ* (see Methods). We derive the former by sampling 100, 000 random iterations of the three parameters of our error model: the true proportion of CpG mutations (from a binomial distribution with *n* = 4933 and *p* = 855/4933, recapitulating the counts from [8]), the proportion of CpG mutations in our data (likewise, with counts of 1165 out of 8108), and the gene conversion adjustment (see above). Taking the mean and standard deviation of this set, we obtain our estimate of *ε* = 1.08 ± 0.28 × 10^−5^. Finally, because of the relatively uniform preparation of our data, the reasonably large standard error, and our limited power to detect statistically significant differences, we use this same error rate for all samples.

### Details of gene conversion correction

As discussed in Methods, the magnitude of the gene conversion effect depends on both the prevalence of non-crossover gene conversion events and the rate at which they introduce heterozygous sites. We make use of a recent direct estimate that non-crossover gene conversion affects 5.9 × 10^−6^ bases per generation [21], which is the product of the rate of events and their average length (on the order of 100 bases in humans [21, 43]). This rate is a genome-wide value, however, and the frequency of gene conversion is highly variable and correlated with local crossover rate [21, 45]. We assume that the gene conversion rate is proportional to the local recombination rate in our genetic map. Because of the way our starting points are chosen, they tend to lie in the relatively recombination-poor regions between hotspots, and so for each of our estimates, we multiply the average 5.9 × 10^−6^ gene conversion rate by the ratio of the local recombination rate (measured in 10 kb windows around the starting points) to the genome-wide average (1.19 cM/Mb, as measured in the deCODE map [16], which was used in [21]). As an example, for our primary eight-genome *H*_5−10_(*d*) curve, the starting-point windows cover 96 Mb but only 51 cM, less than half the average. We note that the differences in average recombination rate per base are only about 10% between the full 100 kb test regions and the central 10 kb windows and 25% or less between the central windows and the full super-regions (S2 Table). To capture noise in the measurement of the local rate, we calculate it separately for each replicate in the jackknife.

The second component of the gene conversion effect is the probability that a gene conversion event introduces a polymorphism that we observe as a heterozygous site. A very simple model for this rate would be that it equals the probability that a randomly chosen base differs from the homologous base on another chromosome in the population (from which it might have been copied in the event of a gene conversion at some point in the past). This quantity can be estimated simply as the heterozygosity in a diploid genome. To this basic model, we add two additional details. First, although our test regions have relatively low heterozygosity (i.e., are relatively recently coalesced), there is also a chance that new mutations that have accumulated on either haplotype could be replaced by the ancestral base via gene conversion. For this probability, we use the heterozygosity at our starting points, *H_S_*(0), divided by 2 (because the new mutations have increased in number since the TMRCA). Second, the probability of mismatch with another chromosome could change over time, and we also find that because of base content, correlated coalescent histories, or other factors, the heterozygosity in other genomes is reduced around our ascertained starting points. For example, other European and East Asian genomes have heterozygosity 5.5–5.9 × 10^−4^ in the 5–10 het regions defined in French B, roughly 15–25% below the genome-wide average (see S1 Table). We assume an average value of 5.7x 10^−4^ both for European test genomes (which have the highest heterozygosity) and other populations, with the exception of Karitiana, for which we instead use their total genome-wide heterozygosity (5.4 × 10^−4^).

Finally, we assign an uncertainty to our correction, which is incorporated into the standard errors we report for our real-data mutation rate estimates. For the rate of gene conversion, we translate the estimated 95% confidence interval of 4.6−7.4 × 10^−6^ per base per generation [21] into a standard deviation of 0.7 × 10^−6^, and for the probability of introducing a new polymorphism, we use a standard deviation of 10% of the total (i.e., 5.7 ± 0.57 × 10^−4^ for most applications).

To test the validity of this approach, we run simulations in which the test data are generated with gene conversion active, and the rest of the procedure is carried out as normal, including the correction just described. For this scenario, we use a true mutation rate of 1.5 × 10^−8^ per base per generation, and for computational efficiency, we assume a uniform rate of gene conversion with *f* = 3.75 (i.e., 3.75 times the average crossover rate) and an average tract length of 100 bases. This results in a gene conversion rate of approximately 4.3 × 10^−6^—about three-quarters of the true rate in humans, but roughly 50% higher than expected for our real-data starting points because of the correlation with local recombination rate. When applying the final correction, we use the average heterozygosity over all simulated genomic segments (approximately 6.6 × 10^−4^) as the “donor” polymorphism probability.

### Details of base content mutability adjustment

Our base content adjustment consists of three elements, which we use to form a ratio of the genome-wide mutation rate to the average local per-base mutation rate around our starting points. First, we compare the fraction of CpG sites between our starting-point regions and the full genome (1.58% after filtering) and use the relative mutation rates from [8], with uncertainty allowed for sampling noise (ratio of *μ*_CpG_/*μ*_other_ = 11.2 ± 0.6). Similarly, we compare total GC content (40% genome-wide), where non-CpG GC bases are assumed to be 50% more mutable than AT bases [44] (including uncertainty, 50±15%). Statistics for our test regions can be found in S2 Table.

Finally, we adjust for the fact that CpG sites are less mutable in regions of high GC content, due to a combination of strand separation effects and low methylation in CpG islands. We modified a result from [46], who estimated an exponential dependence of CpG deamination on local GC content. We (a) used their full CpG transition rates; (b) omitted the tails of the distribution by restricting to GC content between 30% and 60%; and (c) fit a linear dependence instead, both because we are using a genome-wide average and because a linear function appeared to be more accurate in this parameter range. A good fit was provided by the equation *μ*_rel_ = 1 − 2.5Δ_GC_, where *μ*_rel_ is the CpG mutation rate as a proportion of the average (at genome-average GC content of 40%), and A_G_c is the local GC content fraction minus 0.4. Thus, for example, the CpG mutation rate is about 25% higher than average with 30% local GC content and about 50% lower than average with 60% local GC content. We also found empirically from comparing test regions with different CpG and overall GC site frequencies that the coefficient of Δ_GC_ appeared to be approximately 2, in good agreement with the published results. For our final correction, we used a value of 2.5 ± 1.0 for this coefficient.

### Mutation rate heterogeneity

In much of our theoretical and methodological discussion, we have assumed that *μ* is a constant parameter, but in fact different portions of the genome can have different local mutation rates (see Discussion). To learn about the effects that regional variability in *μ* might have on *H_S_*(*d*), we use simulations in which we create data with variability in the mutation rate throughout the genome. Our goal is to approximate the true level of variability present in our data on length scales of several tens of kb, which we do by considering polymorphism rates from populations that are distantly related to those we are using to compute *H_S_*(*d*). Specifically, for each super-region sequence (i.e., each independent segment, including flanking regions; see Methods) being simulated, we divide the length into even thirds and assign a mutation rate for each third proportional to the frequency of doubleton sites (those with exactly two sampled non-reference alleles) in that interval among eight full African genomes (two each San, Dinka, Mbuti, and Yoruba) [24, 28]. The exact rates are chosen in such a way that the overall average rate over the entire genome is equal to a specified value (e.g., 2.5 × 10^−8^). When calling a doubleton, we require at most one individual to be masked (see Methods), and if there are fewer than 10, 000 un-masked sites in any window, we assign the genome-wide average rate for that third of the super-region instead.

We note that while this diversity metric does not precisely capture the true variation in mutation rates across the genome, it should be sufficient for our purposes. The frequency of doubleton sites in the African individuals depends on the local coalescent trees, although this particular statistic should be relatively stable (as opposed to, for example, the total number of polymorphic sites) and will also be smoothed somewhat by considering windows typically on the order of 100 kb. Second, given a local tree, there is additional Poisson noise in the observed counts of doubleton mutations. However, because we are using simulations to gauge the effect of rate variability rather than attempting to add matching variability to the calibration data, we do not need the local estimates to be exact. This is especially true because both sources of stochasticity should cause the apparent variability to be too high rather than too low. Finally, the simulation approach avoids the issue of potential correlations between the African diversity and our non-African data as a result of deep shared ancestry.

Additionally, we note that our estimates should not be impacted by multiple-nucleotide mutation events. It has been estimated that up to a few percent of human point mutations involve nearby clustered nucleotide changes [26, 36, 47], meaning that our inferred rate will correspond to the number of single-base changes per base per generation, rather than the number of independent mutational events per base per generation. However, this is simply a question of how the mutation rate is defined (and moreover, the two are very similar), and all previous mutation rate estimates, to our knowledge, also use the first definition.

### Population heterogeneity and admixture

Our basic model assumes that all of the genomes used to calculate (d) are drawn from the same population. Thus, to the extent that our set of individuals are from groups with different historical sizes, the real data could have different TMRCA patterns in different genomes, whereas the calibration data will be based on the population size profile inferred from the aggregate of all of the samples. A similar phenomenon occurs on a within-genome scale if the test genomes are admixed, in which case individual chromosomes consist of alternating blocks derived from each ancestral mixing population. Although we avoid populations with substantial recent admixture, almost all human populations have experienced some degree of admixture at some point in their history.

Separately, our estimates could potentially be affected by inter-population differences in fine-scale recombination rates. However, the “shared” version of the AA map is intended to apply broadly to non-Africans [17], and it is reasonable to expect that recombination maps will be similar among non-African populations based on their relatively limited diversity of PRDM9 alleles [17, 48, 49].

As with mutation rate heterogeneity, it is difficult to quantify the exact divergence and admixture parameters for the populations under consideration, and hence we use empirical and simulation approaches to study their effects on our inferences. First, while our primary data set consists of a combination of European and East Asian individuals (see Results), we also apply our analyses to the continental groups separately. This allows us both to compare results for populations with different admixture histories and also to compare more homogeneous data sets to the full set with individuals combined from diverged populations.

Second, we perform simulations in which we apply our method to an admixed population designed to emulate present-day Europeans [33, 50]. We simulate a population that experienced a 50/50 mixture 200 generations ago between two populations that had been diverged for 1000 generations. The population sizes are 10, 000 in the shared ancestral population before a bottleneck beginning 2000 generations ago; then 1000 during the bottleneck, which continued until 1000 generations ago (so that the last 200 generations are separate in the two mixing populations); then 15, 000 and 7500 in the two mixing populations after the bottleneck; and finally 25, 000 after the admixture.

### Natural selection

A similar issue to within-genome variation in demography and in the mutation rate is that of heterogeneity in selective effects. Until now we have assumed implicitly that all loci are neutral with respect to fitness, and thus their TMRCAs follow the distributions implied by the standard coalescent model. However, if some sites have non-zero effects on fitness, then two phenomena can result. First, there can be direct selection on new mutations (and also GC-biased gene conversion, which acts similarly to natural selection [21, 44, 51, 52]; see Methods), causing their frequencies to increase or decrease more than expected due to drift. This effect should be minor for our data, since we are considering mutational patterns over relatively short time scales, and if anything, it would likely cause us to underestimate the true mutation rate (see Discussion). The second phenomenon is linked or background selection, whereby local genealogies near selected sites will have different properties from the genome-wide average. In this case, our calibration data, which are based on the average TMRCA distribution, will differ from the real data, but because our PSMC inferences still capture the true genome-wide ancestral population size history, the local variation in this profile will likely only be of second-order importance. Moreover, the effects of linked selection should be similar to those of rate and demographic heterogeneity, since different local genealogical properties caused by selection are analogous to different local mutation rates and/or different local ancestry. Thus, because all three phenomena lead to changes in local levels of diversity, any overall impact of natural selection should be captured in our rate-heterogeneity and admixture simulations described above.

### Symmetry of uncertainty

An implicit assumption we make in computing the uncertainty associated with our estimates is that this uncertainty is symmetric above and below the mean. To test this assumption, we used our results for simulated data. For each of our seven simulation scenarios, we rescaled the 25 independent estimates of *μ* to have mean 0 and standard deviation 1, after which we pooled all 175 data points. The resulting set has a slight positive skewness of 0.14 (see S5 Fig), but this is not statistically significantly different from 0, as the standard deviation of the distribution of sample skewness for 175 observations from a standard normal is approximately 0.18. Thus, we do not see any strong evidence that our uncertainty is greater either above or below the mean.

---

**Figure S1.**
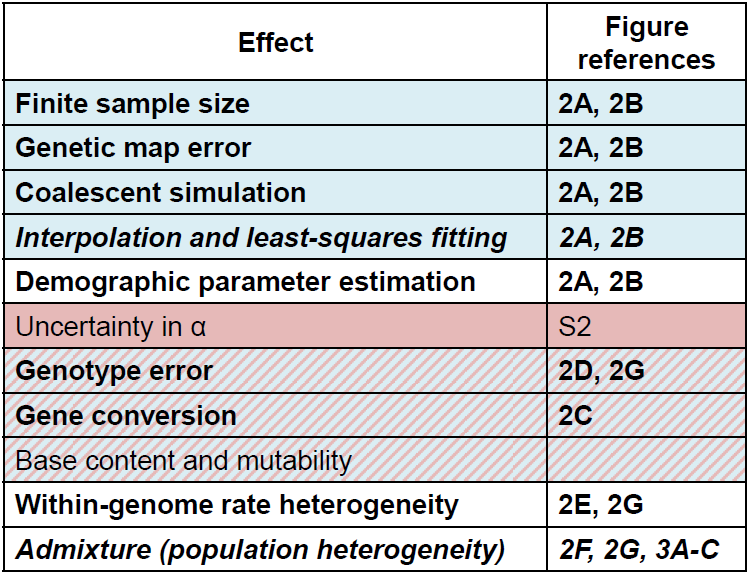
Guide to potential sources of uncertainty associated with our method. Blue shading: included in jackknife procedure; red shading: included in final standard error; cross-hatched shading: uncertainty partially integrated into jackknife and partially included separately in final standard error; bold: tested with simulations; italic: tested empirically with real data. We note that while demographic uncertainty is not explicitly included, we show via simulation that this does not cause our standard error to be underestimated.

**Figure S2.**
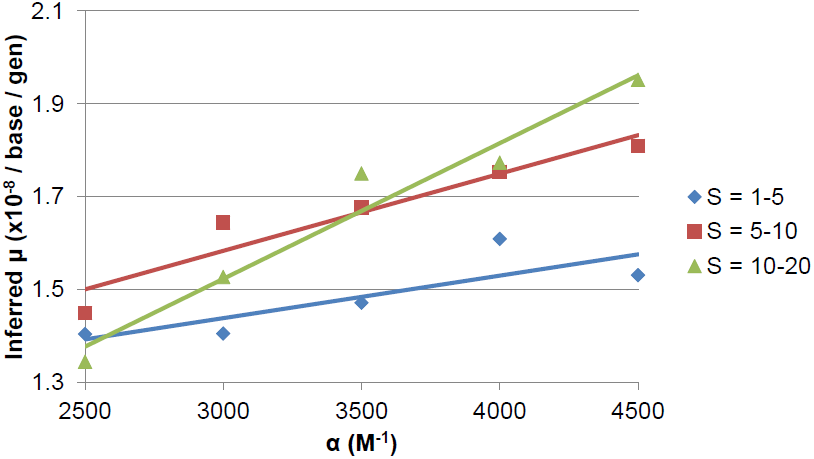
Inferred mutation rates for a range of values of the genetic map error parameter a and starting heterozygosity range *S*. All estimates use our standard data set of eight non-African genomes. Data points represent the inferred rates (independent point estimates), and the lines are linear regression fits for each of the three choices of *S* as a function of *α*. We caution that the values for *S* = 1−5 and 10−20 are less confident than those for our standard range of 5–10. In particular, we believe that the genotype error correction is likely too strong for *S* = 1−5, and thus the 1–5 values here are too low, but we do not have sufficient statistical power to generate a separate error estimate. We also note that the dependence of *μ* on α is stronger for larger *S*, because a higher heterozygosity at the starting points leads to a steeper relaxation of *H_S_*(*d*), so that the curve is more sensitive to the smoothing caused by map error.

**Figure S3.**
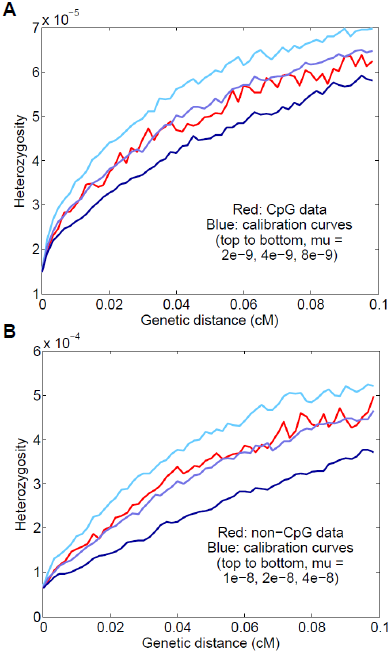
Results for CpG transitions and all other mutations separately, using our primary eight-genome data set. (A) CpGs only; the inferred rate is *μ* = 0.50 ± 0.06 × 10^−8^. (B) Non-CpGs only; the inferred rate is *μ* = 1.36 ± 0.13 × 10^−8^.

**Figure S4.**
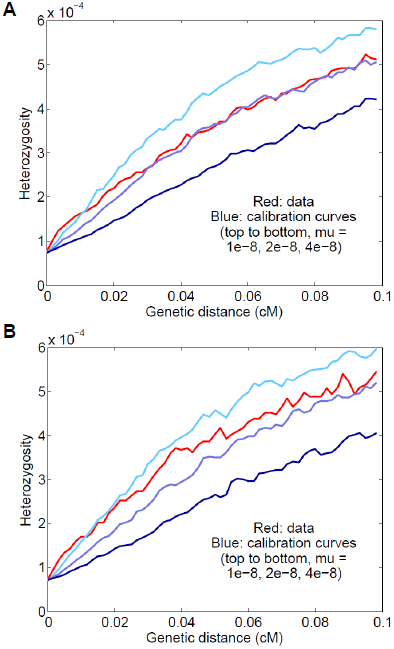
*H*_5−10_(*d*) curves without the pseudo-count prior. (A) Simulated data: we create test data using the prior but omit it for the calibration data. The curve shapes are markedly different, as the calibration curves relax too slowly at the smallest values of *d*. It is also apparent that the inferred value of *μ* is lower than the true value of 2.5 × 10^−8^. (B) Real data for eight non-African genomes. We observe a very similar discrepancy between the real-data and calibration curves (compare Fig 4A).

**Figure S5.**
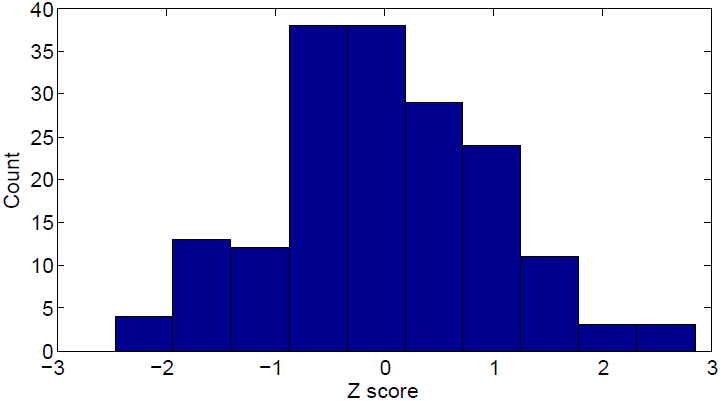
Histogram of Z scores of simulation results. We standardized the 25 independent estimates of *μ* for each of the seven simulated scenarios and combined all 175 values to test for skewness (see S1 Text).

**Table S1.** Sequence divergence for sites passing filters

| Sequence comparison | Per-base diff | Divergence time (My) |
| --- | --- | --- |
| French A | $7.41 \times 10^{-4}$ | $0.67 \pm 0.06$ |
| French B | $7.45 \times 10^{-4}$ | $0.67 \pm 0.06$ |
| Sardinian A | $7.33 \times 10^{-4}$ | $0.66 \pm 0.06$ |
| Sardinian B | $7.23 \times 10^{-4}$ | $0.65 \pm 0.05$ |
| Han A | $7.05 \times 10^{-4}$ | $0.64 \pm 0.05$ |
| Han B | $7.00 \times 10^{-4}$ | $0.63 \pm 0.05$ |
| Dai A | $7.05 \times 10^{-4}$ | $0.63 \pm 0.05$ |
| Dai B | $7.00 \times 10^{-4}$ | $0.63 \pm 0.05$ |
| Australian A | $6.47 \times 10^{-4}$ | $0.58 \pm 0.05$ |
| Australian B | $6.53 \times 10^{-4}$ | $0.59 \pm 0.05$ |
| Karitiana A | $5.35 \times 10^{-4}$ | $0.48 \pm 0.04$ |
| Karitiana B | $5.39 \times 10^{-4}$ | $0.49 \pm 0.04$ |
| Papuan A | $5.95 \times 10^{-4}$ | $0.54 \pm 0.05$ |
| Papuan B | $5.86 \times 10^{-4}$ | $0.53 \pm 0.04$ |
| Human–chimpanzee | $1.23 \times 10^{-2}$ | $11.1 \pm 0.9$ |

Sequence comparisons, with implied divergence times (mean ± standard deviation, in millions of years, using our inferred mutation rate of $1.61 \pm 0.13 \times 10^{-8}$ per base per generation and an average generation interval of 29 years), for sites in the genome passing filters. The first 14 lines represent divergence between the two chromosomes within the individual genomes in our data set (suffix “A” from [24] and suffix “B” from [28], except both Australians from the latter), based on genome-specific filtering (see Methods). Human–chimpanzee statistics are averaged over the filters for the first eight genomes; we note that the third column represents the TMRCA of the two species’ reference sequences rather than the population split time (see Discussion).

**Table S2.**
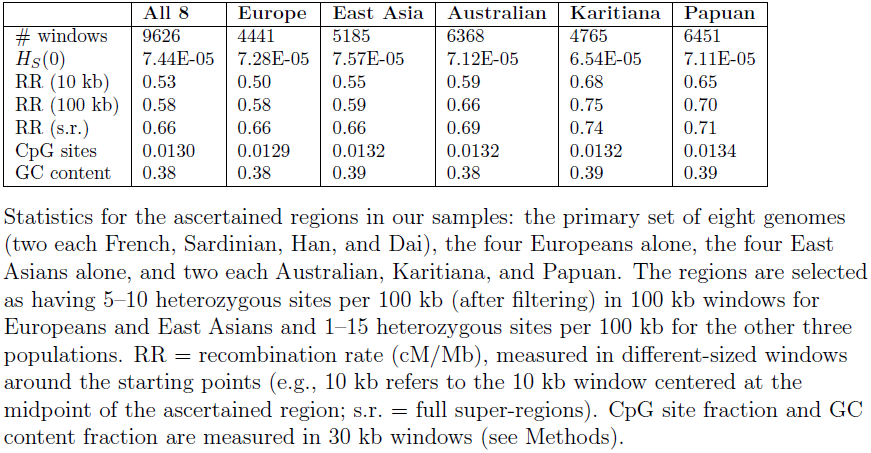
Descriptive statistics for ascertained genomic regions

## References

1. Scally A, Durbin R (2012) Revising the human mutation rate: implications for understanding human evolution. Nat Rev Genet 13: 745–753.

2. Campbell CD, Eichler EE (2013) Properties and rates of germline mutations in humans. Trends Genet 29: 575–584.

3. Ségurel L, Wyman MJ, Przeworski M (2014) Determinants of mutation rate variation in the human germline. Annu Rev Genomics Hum Genet 15: 47–70.

4. Li WH, Tanimura M (1987) The molecular clock runs more slowly in man than in apes and monkeys. Nature 326: 93–96.

5. Kimura M (1968) Evolutionary rate at the molecular level. Nature 217: 624–626.

6. Roach JC, Glusman G, Smit AF, Huff CD, Hubley R, et al. (2010) Analysis of genetic inheritance in a family quartet by whole-genome sequencing. Science 328: 636–639.

7. Conrad DF, Keebler JE, DePristo MA, Lindsay SJ, Zhang Y, et al. (2011) Variation in genome-wide mutation rates within and between human families. Nat Genet 43: 712–714.

8. Kong A, Frigge ML, Masson G, Besenbacher S, Sulem P, et al. (2012) Rate of de novo mutations and the importance of father’s age to disease risk. Nature 488: 471–475.

9. Campbell CD, Chong JX, Malig M, Ko A, Dumont BL, et al. (2012) Estimating the human mutation rate using autozygosity in a founder population. Nat Genet 44: 1277–1281.

10. Besenbacher S, Liu S, Izarzugaza JM, Grove J, Belling K, et al. (2015) Novel variation and de novo mutation rates in population-wide de novo assembled Danish trios. Nat Comm 6: 5969.

11. Fu Q, Li H, Moorjani P, Jay F, Slepchenko SM, et al. (2014) Genome sequence of a 45, 000-year-old modern human from western Siberia. Nature 514: 445–449.

12. Fenner J (2005) Cross-cultural estimation of the human generation interval for use in genetics-based population divergence studies. Am J Phys Anthropol 128: 415–423.

13. Sun JX, Helgason A, Masson G, Ebenesersdottir SS, Li H, et al. (2012) A direct characterization of human mutation based on microsatellites. Nat Genet 44: 1161–1165.

14. Myers S, Bottolo L, Freeman C, McVean G, Donnelly P (2005) A fine-scale map of recombination rates and hotspots across the human genome. Science 310: 321–324.

15. The International HapMap Consortium (2007) A second generation human haplotype map of over 3.1 million SNPs. Nature 449: 851–861.

16. Kong A, Thorleifsson G, Gudbjartsson DF, Masson G, Sigurdsson A, et al. (2010) Fine-scale recombination rate differences between sexes, populations and individuals. Nature 467: 1099–1103.

17. Hinch AG, Tandon A, Patterson N, Song Y, Rohland N, et al. (2011) The landscape of recombination in African Americans. Nature 476: 170–175.

18. Palamara PF, Francioli L, Genovese G, Wilton P, Gusev A, et al. (2015) Leveraging distant relatedness to quantify human mutation and gene conversion rates. Am J Hum Genet doi:10.1016/j.ajhg.2015.10.006.

19. Li H, Durbin R (2011) Inference of human population history from individual whole-genome sequences. Nature 475: 493–496.

20. Sankararaman S, Patterson N, Li H, Pääbo S, Reich D (2012) The date of interbreeding between Neandertals and modern humans. PLoS Genet 8: e1002947.

21. Williams A, Geneovese G, Dyer T, Truax K, Jun G, et al. (2015) Non-crossover gene conversions show strong GC bias and unexpected clustering in humans. eLife 4: e04637.

22. Li H (2014) Towards better understanding of artifacts in variant calling from high-coverage samples. Bioinformatics 30: 2843–2851.

23. Kong A, Thorleifsson G, Frigge ML, Masson G, Gudbjartsson DF, et al. (2014) Common and low-frequency variants associated with genome-wide recombination rate. Nat Genet 46: 11–16.

24. Meyer M, Kircher M, Gansauge MT, Li H, Racimo F, et al. (2012) A high-coverage genome sequence from an archaic Denisovan individual. Science 338: 222–226.

25. Cochran G, Harpending H (2013) Paternal age and genetic load. Hum Biol 85: 515–528.

26. Francioli LC, Polak PP, Koren A, Menelaou A, Chun S, et al. (2015) Genome-wide patterns and properties of de novo mutations in humans. Nat Genet 47: 822–826.

27. Neale BM, Kou Y, Liu L, Maayan A, Samocha KE, et al. (2012) Patterns and rates of exonic de novo mutations in autism spectrum disorders. Nature 485: 242–245.

28. Prüfer K, Racimo F, Patterson N, Jay F, Sankararaman S, et al. (2014) The complete genome sequence of a Neanderthal from the Altai Mountains. Nature 505: 43–49.

29. Genome of the Netherlands Consortium (2014) Whole-genome sequence variation, population structure and demographic history of the Dutch population. Nat Genet 46: 818–825.

30. Mailund T, Halager AE, Westergaard M, Dutheil JY, Munch K, et al. (2012) A new isolation with migration model along complete genomes infers very different divergence processes among closely related great ape species. PLoS Genet 8: e1003125.

31. Prado-Martinez J, Sudmant PH, Kidd JM, Li H, Kelley JL, et al. (2013) Great ape genetic diversity and population history. Nature 499: 471–475.

32. Scally A, Dutheil JY, Hillier LW, Jordan GE, Goodhead I, et al. (2012) Insights into hominid evolution from the gorilla genome sequence. Nature 483: 169–175.

33. Schiffels S, Durbin R (2014) Inferring human population size and separation history from multiple genome sequences. Nat Genet 46: 919–925.

34. Iossifov I, Ronemus M, Levy D, Wang Z, Hakker I, et al. (2012) De novo gene disruptions in children on the autistic spectrum. Neuron 74: 285–299.

35. Fromer M, Pocklington AJ, Kavanagh DH, Williams HJ, Dwyer S, et al. (2014) De novo mutations in schizophrenia implicate synaptic networks. Nature 506: 179–184.

36. Iossifov I, ORoak BJ, Sanders SJ, Ronemus M, Krumm N, et al. (2014) The contribution of de novo coding mutations to autism spectrum disorder. Nature 515: 216–221.

37. Harris K (2015) Evidence for recent, population-specific evolution of the human mutation rate. Proc Natl Acad Sci U S A 112: 3439–3444.

38. McVean GA, Cardin NJ (2005) Approximating the coalescent with recombination. Philos Trans R Soc Lond B Biol Sci 360: 1387–1393.

39. Marjoram P, Wall JD (2006) Fast “coalescent” simulation. BMC Genet 7: 16.

40. Coop G, Wen X, Ober C, Pritchard JK, Przeworski M (2008) High-resolution mapping of crossovers reveals extensive variation in fine-scale recombination patterns among humans. Science 319: 1395–1398.

41. Hellenthal G, Stephens M (2007) mshot: modifying hudson’s ms simulator to incorporate crossover and gene conversion hotspots. Bioinformatics 23: 520–521.

42. Hudson RR (2002) Generating samples under a Wright–Fisher neutral model of genetic variation. Bioinformatics 18: 337–338.

43. Jeffreys AJ, May CA (2004) Intense and highly localized gene conversion activity in human meiotic crossover hot spots. Nat Genet 36: 151–156.

## References

44. Schaibley VM, Zawistowski M, Wegmann D, Ehm MG, Nelson MR, et al. (2013) The influence of genomic context on mutation patterns in the human genome inferred from rare variants. Genome Res 23: 1974–1984.

45. Pratto F, Brick K, Khil P, Smagulova F, Petukhova GV, et al. (2014) Recombination initiation maps of individual human genomes. Science 346: 1256442.

46. Fryxell KJ, Moon WJ (2005) CpG mutation rates in the human genome are highly dependent on local GC content. Mol Biol Evol 22: 650–658.

47. Harris K, Nielsen R (2014) Error-prone polymerase activity causes multinucleotide mutations in humans. Genome Res 24: 1445–1454.

48. Baudat F, Buard J, Grey C, Fledel-Alon A, Ober C, et al. (2010) PRDM9 is a major determinant of meiotic recombination hotspots in humans and mice. Science 327: 836–840.

49. Berg IL, Neumann R, Lam KWG, Sarbajna S, Odenthal-Hesse L, et al. (2010) PRDM9 variation strongly influences recombination hot-spot activity and meiotic instability in humans. Nat Genet 42: 859–863.

50. Lazaridis I, Patterson N, Mittnik A, Renaud G, Mallick S, et al. (2014) Ancient human genomes suggest three ancestral populations for present-day Europeans. Nature 513: 409–413.

51. Capra JA, Hubisz MJ, Kostka D, Pollard KS, Siepel A (2013) A model-based analysis of GC-biased gene conversion in the human and chimpanzee genomes. PLoS Genet 9: e1003684.

52. Lachance J, Tishkoff SA (2014) Biased gene conversion skews allele frequencies in human populations, increasing the disease burden of recessive alleles. Am J Hum Genet 95: 408–420.

